# A PARP14/TARG1-Regulated RACK1 MARylation Cycle Drives Stress Granule Dynamics in Ovarian Cancer Cells

**DOI:** 10.1101/2023.10.13.562273

**Authors:** Sridevi Challa, Tulip Nandu, Hyung Bum Kim, Xuan Gong, Charles W. Renshaw, Wan-Chen Li, Xinrui Tan, Marwa W. Aljardali, Cristel V. Camacho, Jin Chen, W. Lee Kraus

## Abstract

Mono(ADP-ribosyl)ation (MARylation) is emerging as a critical regulator of ribosome function and translation. Herein, we demonstrate that RACK1, an integral component of the ribosome, is MARylated on three acidic residues by the mono(ADP-ribosyl) transferase (MART) PARP14 in ovarian cancer cells. MARylation of RACK1 is required for stress granule formation and promotes the colocalization of RACK1 in stress granules with G3BP1, eIF3η, and 40S ribosomal proteins. In parallel, we observed reduced translation of a subset of mRNAs, including those encoding key cancer regulators (e.g., AKT). Treatment with a PARP14 inhibitor or mutation of the sites of MARylation on RACK1 blocks these outcomes, as well as the growth of ovarian cancer cells in culture and in vivo. To re-set the system after prolonged stress and recovery, the ADP-ribosyl hydrolase TARG1 deMARylates RACK1, leading to the dissociation of the stress granules and the restoration of translation. Collectively, our results demonstrate a therapeutically targetable pathway that controls stress granule assembly and disassembly in ovarian cancer cells.

**Summary:** We have discovered a druggable PARP14/TARG1-regulated pathway that mediates site- specific mono(ADP-ribosyl)ation of RACK1, a ribosomal protein. This pathway controls stress granule assembly and disassembly, as well as the translation of a subset of mRNAs, to modulate the growth of ovarian cancer cells in culture and in vivo.

## Introduction

The properly controlled translation of mRNAs by ribosomes is fundamentally important to cellular functions and broader biological outcomes (Klinge and Woolford, 2019; Wu et al., 2020). A growing body of work has linked ribosome biogenesis, ribosome function, and translation to cellular outcomes in cancers (Brighenti et al., 2015; Bustelo and Dosil, 2017). The central components of the ribosome, including the repertoire of ribosomal proteins, can be regulated and diversified to control protein translation (Sauert et al., 2015). This regulation is mediated, in part, by post-translational modifications (PTMs) of ribosomal proteins, including phosphorylation and ubiquitylation, among others (Simsek and Barna, 2017). Anticancer therapies that block ribosomal function induce the assembly of stress granules (Zhou et al., 2023). Stress granule assembly is a crucial mechanism that cells use to coordinate selective translation of transcripts vital for cell survival by compartmentalizing mRNAs, translation machinery, and apoptotic signaling proteins (Park et al., 2020).

We recently identified sites of mono(ADP-ribosyl)ation (MARylation) on a number of ribosomal proteins in mammalian cells (Challa et al., 2021a). MARylation is a reversible PTM of proteins catalyzed by mono(ADP-ribosyl) transferases (MARTs), resulting in the covalent attachment of a single ADP-ribose (ADPR) moiety derived from NAD^+^ on a variety of amino acid residues (*e.g.*, Glu, Asp, Ser) (Challa et al., 2021b; Gibson and Kraus, 2012; Schreiber et al., 2006). Although the role of MARylation in cancer is not well understood, the roles of MARylation in cellular stress responses to viral and bacterial infection are well characterized (Challa et al., 2021b; Kim et al., 2020). Recent studies have begun to reveal novel and interesting functions for cytoplasmic MARTs, such as PARP7, PARP12, PARP14, and PARP16, in molecular and cellular functions including RNA processing, translation, stress granule formation, unfolded protein response, and regulation of the cytoskeleton (Ahmed et al., 2015; Bindesboll et al., 2016; Challa et al., 2021a; Di Paola et al., 2012; Iwata et al., 2016; Jwa and Chang, 2012; Leung et al., 2011; Palavalli Parsons et al., 2021; Roper et al., 2014; Vyas et al., 2013; Vyas et al., 2014).

In a recent study, we demonstrated that two specific sets of ribosomal proteins are MARylated: (1) “assembly factors” (e.g., RPS6, RPL24) located at the interface between the 60S and 40S ribosomal subunits and (2) “regulatory factors” (e.g., receptor for activated C kinase 1, RACK1) located on the surface of the 40S subunit (Challa et al., 2021a). These different ribosome MARylation events may have distinct functions in regulating ribosome biogenesis and ribosome function, respectively. In ovarian cancer cells, PARP16 uses NAD^+^ produced by the cytosolic NAD^+^ synthase NMNAT-2 to MARylate RPS6 and RPL24 (Challa et al., 2021a). The MARylation attenuates translation to help maintain proteostasis and promote the growth of the cancer cells. The function of RACK1 MARylation and the MART that mediates it have not been characterized.

RACK1 is an integral ribosome component (Rabl et al., 2011) and member of the tryptophan-aspartate repeat (WD-repeat) family of proteins (Adams et al., 2011). It serves as chaperone that shuttles proteins around the cell and anchors them where needed (Adams et al., 2011). In addition, it plays key roles in cancers (Li and Xie, 2015) and has been linked to stress granule formation (Buchan and Parker, 2009; Zhou et al., 2023). Stress granules are inducible, membrane-less condensates enriched in mRNAs, RNA-binding proteins, and 40S ribosomal subunits (Asadi et al., 2021; Zhou et al., 2023). Stress granule assembly, which can be tracked by G3BP1, a marker of stress granule formation (Asadi et al., 2021; Zhou et al., 2023), is an adaptive mechanism in response to cellular stressors, which results in reduced translation until the stress can be removed (Asadi et al., 2021; Zhou et al., 2023).

The therapeutic potential of ADP-ribosyltransferases has received considerable attention due to the U.S. FDA’s approval of the use of four different PARP inhibitors for the treatment of ovarian and breast cancers (Bitler et al., 2017; Clamp and Jayson, 2015; Liu and Matulonis, 2016). Beyond these drugs, which target nuclear enzymes, there is a growing interest in drugging cytosolic MARTs (Peng et al., 2017; Yoneyama-Hirozane et al., 2017). In fact, a number of academic labs and pharmaceutical companies are developing chemical inhibitors to tap the unexplored therapeutic potential of MARTs, including inhibitors of PARP7 (Gozgit et al., 2021; Sanderson et al., 2023), PARP14 (Schenkel et al., 2021), and PARP16 (Bejan et al., 2022). Our current studies aim to characterize additional mechanisms through which ribosome MARylation regulates stress responses in cancer cells, with a focus on RACKl-mediated regulation of stress granule formation and the control of ribosome function.

## Results

### RACK1, an integral component of the 40S ribosomal subunit, is MARylated in ovarian cancer cells

Our recent studies have demonstrated the control of ribosome function by MARylation, a PTM resulting in the covalent attachment of ADP-ribose by MART enzymes (Challa et al., 2021a). RACK1 is an integral component of the ribosome (Rabl et al., 2011) located in a regulatory region of the 40S subunit (Park et al., 2020). It functions as a scaffolding protein, which recruits proteins that are important for quality control during mRNA translation (Arimoto et al., 2008), and serves as an essential component of stress granules (Buchan and Parker, 2009; Zhou et al., 2023). We previously identified sites of MARylation on RACK1 in ovarian cancer cells using mass spectrometry-based proteomics (Challa et al., 2021a) (Fig. 1A; Asp 144, Glu 145, and Asp 203). Herein, we confirmed that RACK1 is MARylated in OVCAR3 ovarian cancer cells using immunoprecipitation of endogenous RACK1, followed by immunoblotting with an antibody-like MAR detection reagent (Gibson et al., 2017a) (Fig. 1B; Suppl. Fig. S1, A and B). Next, we generated OVCAR3 cells that, when cultured in doxycycline (Dox), simultaneously knockdown endogenous RACK1 and ectopically express HA-tagged wild-type RACK1 (RACK1-WT) or RACK1 with all the three MAR acceptor sites mutated. We refer to the RACK1 protein with the three MAR acceptor sites mutated (i.e., D144N, E145Q, D203N) as “RACK1-Mut”. Immunoprecipitation of the HA-tagged RACK1 followed by immunoblotting for MAR demonstrated site-specific MARylation of RACK1 at Asp 144, Glu 145, and Asp 203 (Fig. 1C). Generating antibodies to detect site-specific MARylation is technically challenging due to limitations in the synthesis of site-specific MARylated antigens. To overcome this limitation, we developed a proximity ligation assay (PLA) for *in situ* detection of site-specific MARylation using MAR and HA (i.e., RACK1) antibodies, which confirmed robust MARylation of RACK1-WT, with significantly reduced MARylation of mutant RACK1 (Fig. 1, D and E).

**Figure 1.**
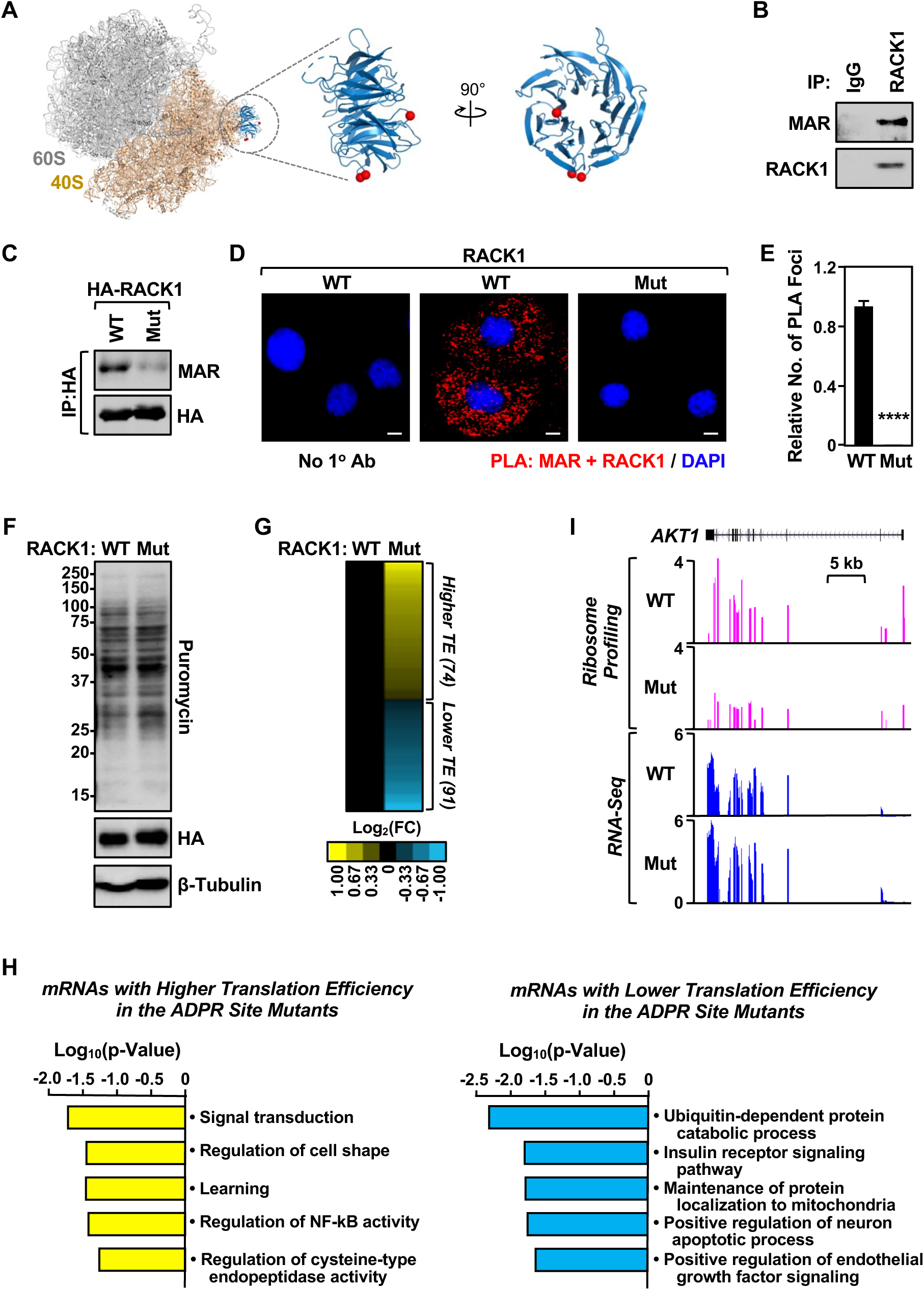
Site-specific MARylation of RACK1 in ovarian cancer cells. **(A)** *(Left)* Spatial distribution of the proteins modified by MARylation in the 80S ribosome (PDB ID: 4V6X). (*Middle and right*) Sites of MARylation within RACK1 (Asp 144, Glu 145, and Asp 203; blue ribbon) are indicated in two expanded views, with the structure in the right rotated by 90°. **(B)** RACK1 is MARylated. Endogenous RACK1 was immunoprecipitated from OVCAR3 cells and subjected to immunoblotting for MAR and RACK1. **(C)** RACK1 is MARylated at Asp 144, Glu 145, and Asp 203. HA-tagged RACK1 was immunoprecipitated from OVCAR3 cells ectopically expressing wild-type (WT) or MARylation site mutant (Mut) RACK1 and subjected to immunoblotting for MAR and HA. **(D)** In situ detection of RACK1 MARylation. Proximity ligation assay (PLA) of RACK1 and MAR in OVCAR3 cells subjected to Dox-induced knockdown of endogenous and re-expression of RACK1 (WT or Mut). DNA was stained with DAPI. Scale bar is 15 µm. **(E)** Quantification of multiple experiments like the one shown in panel (D). Each bar represents the mean + SEM of MAR-RACK1 PLA foci from three biological replicates (Student’s t-test, **** p < 0.0001). **(F)** RACK1-Mut expression does not alter global protein synthesis in OVCAR3 cells. Immunoblot analysis of puromycin incorporation assays from OVCAR3 cells subjected to Dox- induced knockdown of endogenous and re-expression of RACK1. β-tubulin serves as a loading control. **(G and H)** Regulation of mRNA translation by RACK1 MARylation. Ribosome profiling of OVCAR3 cells subjected to Dox-induced knockdown of endogenous RACK1 followed by re- expression of exogenous RACK1 (WT or Mut). (G) Heatmap representation of mRNAs that exhibit altered translation efficiency when RACK1-Mut was expressed. (H) Gene ontology enrichment analysis of the translationally upregulated and downregulated mRNAs. **(I)** RACK1 MARylation regulates translation of *AKT1*. Example ribosome profiling and RNA- seq traces of *AKT1* in OVCAR3 cells subjected to Dox-induced knockdown of endogenous RACK1 and re-expression of exogenous RACK1 (WT or Mut). A schematic of the *AKT1* gene with a scale bar is shown.

### RACK1 is required for efficient translation of selected mRNAs

Our previous studies demonstrated that one function of MARylation of ribosomal proteins is to inhibit global protein synthesis by altering polysome formation. Therefore, we measured global protein synthesis levels of RACK1-WT- or RACK1-Mut-expressing cells using puromycin incorporation assays. We did not observe an obvious change in global protein synthesis in cells deficient in RACK1 MARylation (Fig. 1F). We next performed ribosome profiling (Ribo-seq) assays (Chen et al., 2020; McGlincy and Ingolia, 2017) to investigate potential changes in translational efficiency in cells deficient in RACK1 MARylation. We observed changes in the translation levels of 165 transcripts in RACK1-Mut-expressing cells versus RACK1-WT-expressing cells (Fig. 1G). Gene ontology of the affected transcripts (up or down) showed enrichment in mRNAs encoding proteins involved in receptor tyrosine kinase signaling, including *AKT1* (Fig. 1, H and I). These results implicate RACK1 MARylation in the control of translation of a subset of mRNAs.

### Site-specific MARylation of RACK1 is required for stress granule assembly

Our results demonstrate that RACK1 controls the translation of proteins that play an important role in ovarian cancer biology, although the mechanisms through which translation is regulated remains unknown. Since RACK1 is a key player in stress granule assembly (Buchan and Parker, 2009; Zhou et al., 2023), we tested whether MARylation alters the localization of RACK1 to stress granules. Co-immunoprecipitation assays and PLAs in OVCAR3 cells expressing RACK1-WT or RACK1-Mut show that MARylation of RACK1 is required for its interaction with G3BP1, a key nucleating factor for stress granules (Marcelo et al., 2021) (Fig. 2, A and B; Suppl. Fig. S1, C and D). We next performed polysome profiling to investigate the effect of RACK1 MARylation on its incorporation into ribosomes. Interestingly, while RACK1 MARylation was not required for the integration of RACK1 into ribosomes, loss of RACK1 MARylation by mutation of the sites inhibited the association of G3BP1 with ribosomes (Fig. 2, C and D; see the reduced levels of G3BP1 in fractions 6-15). These results demonstrate that loss of RACK1 MARylation reduces the interaction of G3BP1 with ribosomes.

**Figure 2.**
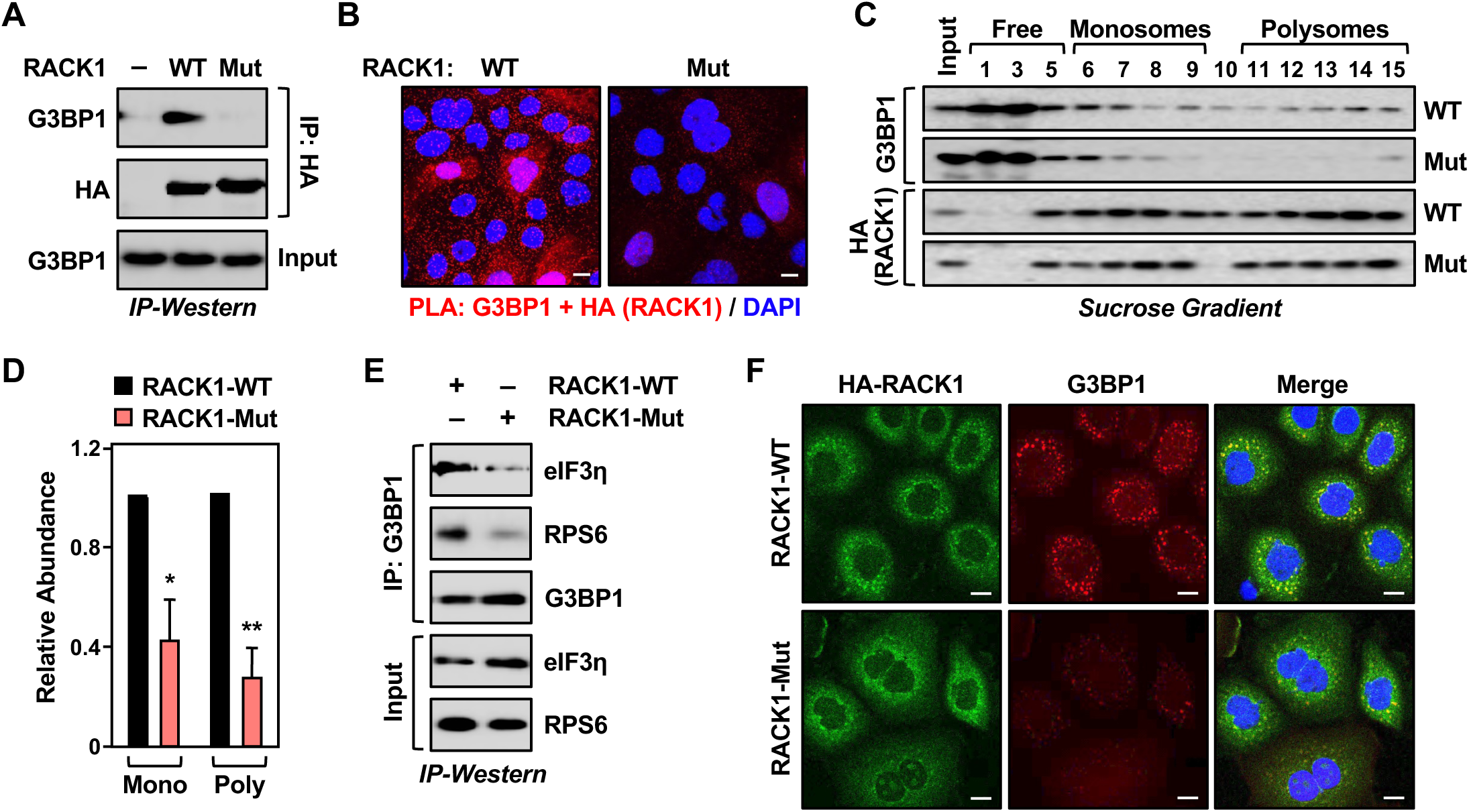
Site-specific MARylation of RACK1 is required for stress granule assembly. **(A and B)** Loss of RACK1 MARylation inhibits RACK1 interaction with G3BP1. (A) HA- tagged RACK1 was immunoprecipitated from OVCAR3 cells with Dox-induced knockdown of endogenous RACK1 and re-expression of exogenous RACK1. The immunoprecipitates were subjected to immunoblotting for G3BP1 and HA. (B) PLA using G3BP1 and HA antibodies. DNA was stained with DAPI. Scale bar is 15 µm. **(C and D)** Loss of RACK1 MARylation inhibits the recruitment of G3BP1 to ribosomes. (C) Immunoblot analysis for HA-tagged RACK1 and G3BP1 in sucrose density gradient fractions of ribosomes prepared from OVCAR3 cells subjected to Dox-induced knockdown of endogenous RACK1 and re-expression of exogenous RACK1. Each bar in the graph in (D) represents the mean + SEM of the relative abundance of G3BP1 in monosomes or polysomes (n = 3, two-way ANOVA, * p < 0.05 and ** p<0.01). **(E and F)** Loss of RACK1 MARylation inhibits G3BP1 localization to stress granules and its interaction with translation factors that are key components of stress granules. (E) G3BP1 was immunoprecipitated from OVCAR3 cells with Dox-induced knockdown of endogenous RACK1 and re-expression of exogenous RACK1. The immunoprecipitates were subjected to immunoblotting for eIF3η, RPS6, and G3BP1 as indicated. (F) Immunofluorescent staining assays of OVCAR3 cells with Dox-induced knockdown of endogenous RACK1 and re- expression of exogenous RACK1 subjected to 15 minutes of treatment with 250 µM sodium arsenite (NaAsO_2_). Staining for HA (RACK1) and G3BP1. DNA was stained with DAPI. Scale bar is 15 µm.

We confirmed this observation using co-immunoprecipitation analysis of G3BP1- interacting proteins in OVCAR3 cells. We observed that in cells expressing RACK1-Mut, G3BP1 failed to interact with eIF3η, another stress granule marker protein (Zhou et al., 2023), and RPS6, a component of the 40S ribosomal subunit (Fig. 2E; Suppl. Fig. S1E). These results demonstrate that RACK1-MARylation drives the association of G3BP1 with ribosomes. Because G3BP1 is the central nucleating factor for stress granule assembly, we determined how the loss of RACK1 MARylation might impact granule assembly. Immunofluorescence staining assays in OVCAR3 cells expressing RACK1-Mut exhibited reduced levels of G3BP1 puncta compared to cells expressing RACK1-WT (Fig. 2F; Suppl. Fig. S1F). These assays also demonstrated that RACK1-Mut cells have reduced localization of RACK1, RPS6, and eIF3η to stress granules (Fig. 2F; Suppl. Fig. S1, G and H). Treating the cells with puromycin (15 or 30 minutes), which destabilizes polysome formation (Kedersha et al., 2000), normalized stress granule assembly between NaAsO_2_-treated cells expressing RACK1-WT and those expressing RACK1-Mut (Suppl. Fig. S1, I through L). We confirmed that MARylation does not play a role in translation repression, as treatment with puromycin under stress conditions (i.e., +NaAsO_2_) did not affect translation as assessed by puromycin incorporation (Suppl. Fig. S1, M and N). Collectively, these data demonstrate that (1) RACK1 MARylation regulates stress granule assembly by regulating polysome function and (2) site-specific MARylation of RACK1 drives protein-protein interactions that are required for stress granule assembly.

### PARP14 inhibition reduces stress granule assembly

In our previous work, we identified PARP16 as a MART that MARylates selected ribosomal proteins to regulate the loading of mRNAs onto ribosomes and their translation (Challa et al., 2021a). To determine which MART MARylates RACK1 in OVCAR3 cells, we used the RACK1+MAR PLA coupled with an siRNA screen of MART enzymes, focusing on those that are both expressed in OVCAR3 cells and are primarily cytosolic. We observed that knockdown of *PARP14* mRNA caused the most consistent and dramatic reduction in RACK1 MARylation (Suppl. Fig. S2A). PARP14 is a macrodomain-containing MART that has been implicated in stress responses and cancer (Dhoonmoon and Nicolae, 2023; Dukic et al., 2023; Torretta et al., 2023).

To confirm the role of PARP14 in the MARylation of RACK1, we used a chemical inhibitor of PARP14 (PARP14i), RBN012579 (Schenkel et al., 2021), which inhibits PARP14 activity, as shown by a reduction in autoMARylation (Fig. 3A). Treatment with PARP14i also inhibited RACK1 MARylation in a PLA (Fig. 3B; Suppl. Fig. S2B). We also observed that chemical inhibition of PARP14 activity phenocopies the expression of RACK1-Mut in various endpoint assays: (1) reduced association of G3BP1 with ribosomes (Fig. 3, C and D), (3) reduced interactions between G3BP1 and stress granule factors (Fig. 3, E through H), and (3) reduced stress granule assembly (Fig. 3, I and J; Suppl. Fig. S2, C and D). We observed similar regulation of PARP14-mediated RACK1-MARylation and stress granule assembly in additional ovarian cancer cell lines, SKOV3 and HCC5044 (Suppl. Fig. S2, E through H; note the reduced number of discrete G3BP1 puncta upon PARP14i treatment). Together, these data show that PARP14-mediated, site-specific MARylation of RACK1 drives stress granule assembly in ovarian cancer cells.

**Figure 3.**
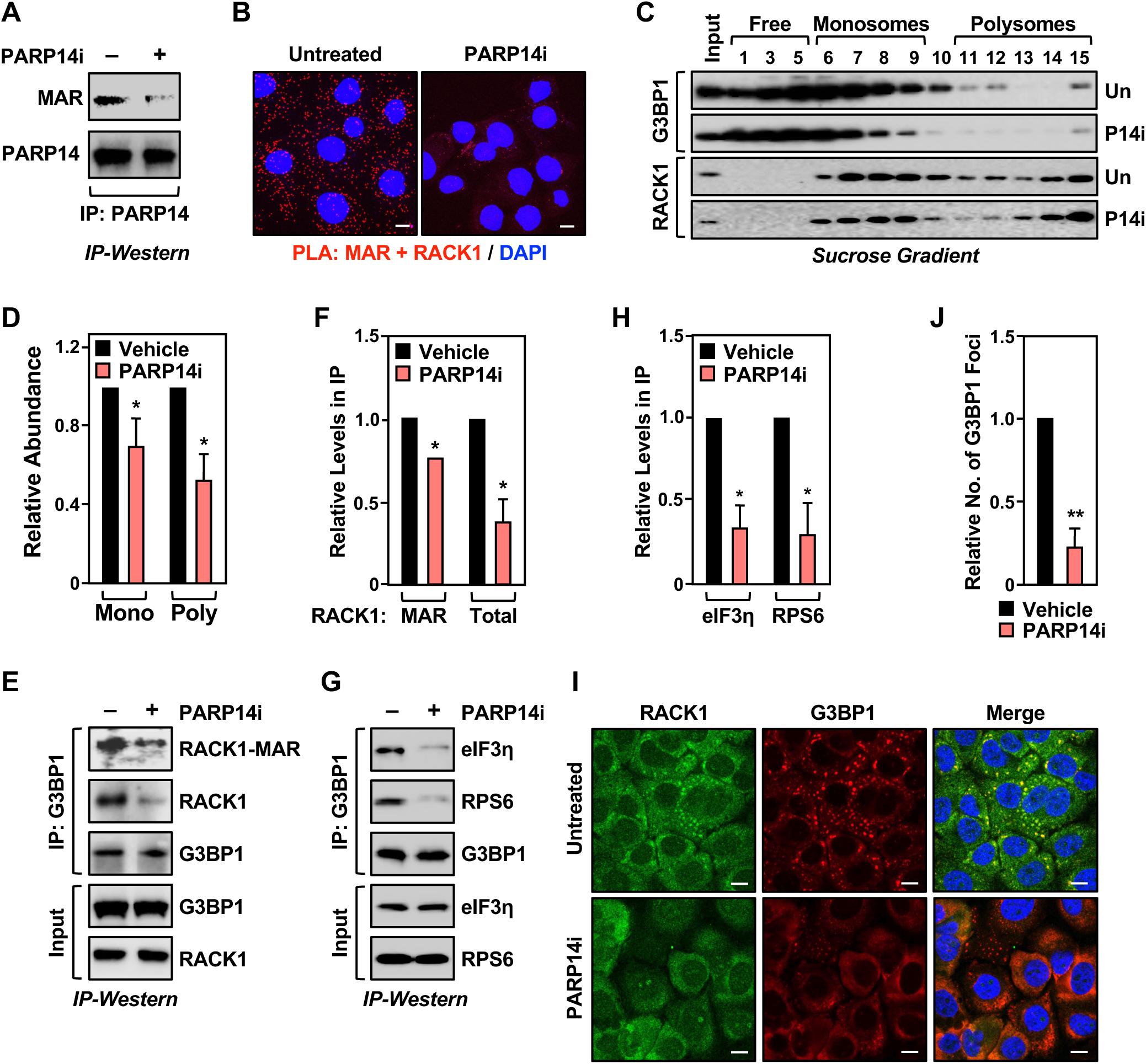
PARP14 inhibition reduces stress granule assembly. **(A)** PARP14 inhibitor blocks PARP14 autoMARylation. OVCAR3 cells were treated with 10 µM PARP14 inhibitor (RBN012759) for 24 hours. PARP14 was immunoprecipitated and subjected to immunoblotting for PARP14 and MAR. **(B)** Inhibition of PARP14 catalytic activity blocks RACK1 MARylation. PLA using MAR and RACK1 antibodies in OVCAR3 cells treated with 10 µM PARP14 inhibitor (RBN012759) for 24 hours. DNA was stained with DAPI. Scale bar is 15 µm. **(C and D)** PARP14 inhibition reduces the recruitment of G3BP1 to ribosomes. (C) Immunoblot analysis of RACK1 and G3BP1 in sucrose density gradient fractions of ribosomes prepared from OVCAR3 cells treated with 10 µM PARP14 inhibitor for 24 hours. Each bar in the graph in (D) represents the mean + SEM of the relative abundance of G3BP1 in monosomes or polysomes (n = 3, two-way ANOVA, * p < 0.05 and ** p<0.01). **(E and F)** PARP14 inhibition reduces G3BP1 interaction with RACK1. (E) G3BP1 was immunoprecipitated from OVCAR3 cells treated with 10 µM PARP14 inhibitor for 24 hours and subjected to immunoblotting for MAR, RACK1, and G3BP1 as indicated. The band corresponding to the molecular weight of RACK1 was indicated as MARylated RACK1. Each bar in the graph in (F) represents the mean + SEM of the relative abundance of total RACK1 or MARylated RACK1 in G3BP1 immunoprecipitates (n = 3, Student’s t-test, * p < 0.05). **(G and H)** PARP14 inhibition reduces G3BP1 interaction with translation factors that are key components of stress granules. (G) G3BP1 was immunoprecipitated from OVCAR3 cells treated with 10 µM PARP14 inhibitor for 24 hours and subjected to immunoblotting for eIF3η, RPS6, and G3BP1 as indicated. Each bar in the graph in (H) represents the mean + SEM of the relative abundance of eIF3η and RPS6 in G3BP1 immunoprecipitates (n = 3, Student’s t-test, * p < 0.05). **(I and J)** PARP14 inhibition reduces G3BP1 localization to stress granules. Immunofluorescent staining assays of OVCAR3 cells treated with 10 µM PARP14 inhibitor for 24 hours and subjected to 15 minutes of treatment with 250 µM sodium arsenite (NaAsO_2_). Staining for RACK1 and G3BP1. DNA was stained with DAPI. Scale bar is 15 µm. Each bar in the graph in (J) represents the mean + SEM of the relative abundance of stress granules (n = 3, Student’s t- test, ** p<0.01).

### Loss of RACK1 MARylation sensitizes ovarian cancer cells to stress

Since loss of RACK1-MARylation suppresses stress granule assembly, which is crucial for overcoming stress (Arimoto et al., 2008; Park et al., 2020), we surmised that the loss of PARP14-mediated site-specific MARylation of RACK1 will sensitize the cells to external stressors. To test this, we performed cell growth assays using ovarian cancer cells cultured in the presence of thapsigargin, which induces endoplasmic reticulum (ER) stress (Sagara and Inesi, 1991), as well as carboplatin, which induces oxidative stress (Yu et al., 2018). Ovarian cancer cells expressing RACK1-Mut or treated with PARP14i exhibited slower growth than cells expressing RACK1-WT or treated with vehicle in the presence of thapsigargin or carboplatin (Fig. 4, A and B; Suppl. Fig. S3, A through D). Similar effects were observed in OVCAR3 xenograft tumors grown in immunodeficient mice (Fig. 4, C and D; Suppl. Fig. S3, E through G). Further analysis demonstrated that the expression of RACK1-Mut or PARP14i treatment increased ER stress as indicated by increased phosphorylation of eIF2a, causing apoptosis as indicated by increased cleaved caspase-3 (Suppl. Fig. S3, H through K). These results connect the site-specific MARylation of RACK1 to cellular and biological outcomes. In addition, they provide additional evidence for biological connections between ER stress and stress granules (Nicchitta, 2022).

**Figure 4.**
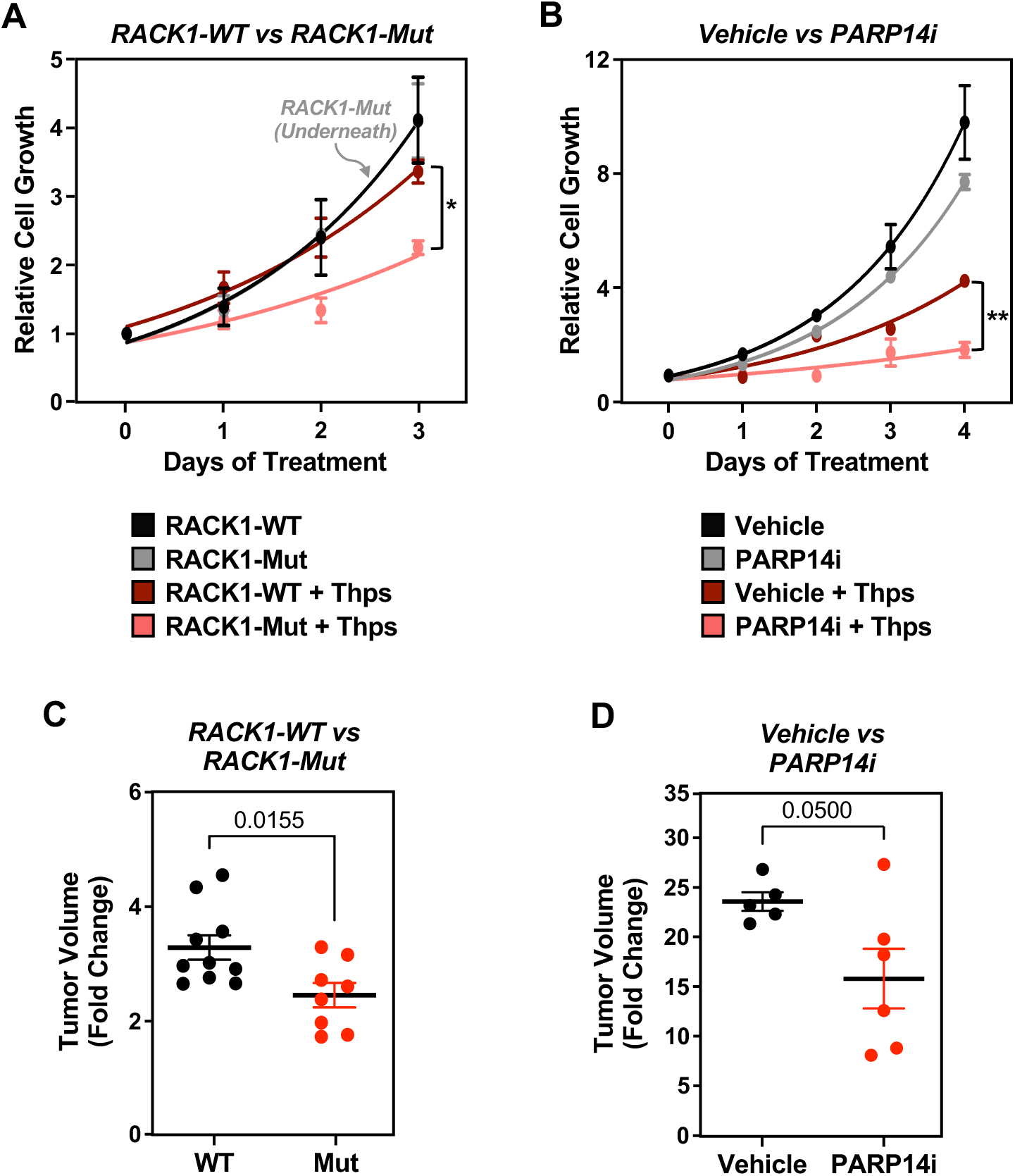
Loss of RACK1 MARylation sensitizes ovarian cancer cells to stress and inhibits their growth. **(A)** RACK1-Mut expressing cells are sensitive to ER stress, which inhibits their growth. Growth curves of OVCAR3 cells with Dox-induced knockdown of endogenous RACK1 and re-expression of exogenous RACK1 (WT or Mut) in the presence or absence of 3 nM thapsigargin (Thps) for the indicated times. The arrow points to the RACK-Mut growth curve beneath the RACK-WT growth curve under basal conditions. Each point represents the mean ± SEM of the growth of the cells relative to Day 0 of treatment (n = 3, two-way ANOVA, * p < 0.01). **(B)** PARP14 inhibition sensitizes ovarian cancer cells to ER stress and inhibits their growth. Growth curves of OVCAR3 cells in the presence or absence of 10 µM PARP14 inhibitor (PARP14i) and 3 nM thapsigargin (Thps) for the indicated times. Each point represents the mean ± SEM of the growth of the cells relative to Day 0 of treatments (n = 3, two-way ANOVA, ** p < 0.001). **(C and D)** Expression of RACK1-Mut or treatment with PARP14 inhibitor (PARP14i) inhibits the growth of OVCAR3 xenograft tumors derived from cells like those described in (A) and (B). The xenograft tumors were established in immunocompromised NSG mice subjected to the treatments indicated and grown until the mice reached the end-point for euthanasia as required by IACUC. (C) Tumor volume at Day 69. Each cluster in the graph shows the mean and the individual data points for n = 10 or 8 mice (WT or Mut, respectively), Student’s t-test, p = 0.0155. (D) Tumor volume at Day 19 post-treatment. Each cluster in the graph shows the mean and the individual data points for n = 5 or 6 mice (vehicle or PARP14i, respectively), Student’s t-test, p = 0.05. Different timelines in the two xenograft experiments were dictated by different growth rates of parental (D) versus Dox-treated cells (C).

### Loss of TARG1 enhances stress granule assembly by increasing RACK1 MARylation

Our studies thus far have demonstrated that site-specific MARylation of RACK1 mediated by PARP14 is required for stress granule assembly and stress responses in ovarian cancer cells. Stress granule assembly is a dynamic process; the coordinated regulation of stress granule assembly and disassembly drives mRNA localization and translational regulation. Although our results identified a PARP14- and RACK1-dependent pathway of stress granule assembly, the mechanisms that control disassembly of stress granules in this pathway are unknown. We considered the possibility that the removal of RACK1 MARylation by an ADP-ribosyl hydrolase might serve this role.

To test this, we performed an siRNA screen using siRNAs targeting known ADP-ribosyl hydrolases (Suppl. Fig. S4A), which revealed that knockdown of *TARG1* mRNA (from the *OARD1* gene) dramatically increased the MARylation of ribosomal proteins (Suppl. Fig. S4B). TARG1 is an ADP-ribosyl hydrolase that can specifically remove the terminal ADPR on Glu and Asp residues (Sharifi et al., 2013). Recent studies have demonstrated that TARG1 interacts with ribosomal proteins and localizes to stress granules (Butepage et al., 2018; Zaja et al., 2020). The results from our screen and prior studies inspired us to investigate the functional role of TARG1 in stress granule assembly in more detail.

In this regard, we observed that depletion of TARG1 increased RACK1 MARylation, both by IP-Western and PLAs (Fig. 5, A and B; Suppl Fig. 5, A and B) and stress granule assembly (Fig. 5, C and D; Suppl. Fig S5C) in OVCAR3 cells, as well as two other ovarian cancer cell lines (Suppl Fig. 5, D through G). Since our initial results showed that site-specific MARylation of RACK1 controls the translation of mRNAs that are important for the survival of ovarian cancer cells (i.e., *AKT1*) (Fig. 1, F through I), we sought to explore the possible role of TARG1 in the regulation of translation. We performed ribosome profiling assays using OVCAR3 cells subjected to siRNA-mediated knockdown of *TARG1/OARD1*. The results demonstrated that depletion of TARG1 led to an increase in the translation of mRNAs encoding proteins involved in DNA repair and DNA replication (Fig. 5E). Interestingly, depletion of TARG1 also led to a decrease in the translation and transcription of mRNAs encoding proteins involved in translation (Fig. 5E). Collectively, these results suggest a role for TARG1 in controlling a coordinated translation program in response to cellular stress via RACK1 deMARylation.

**Figure 5.**
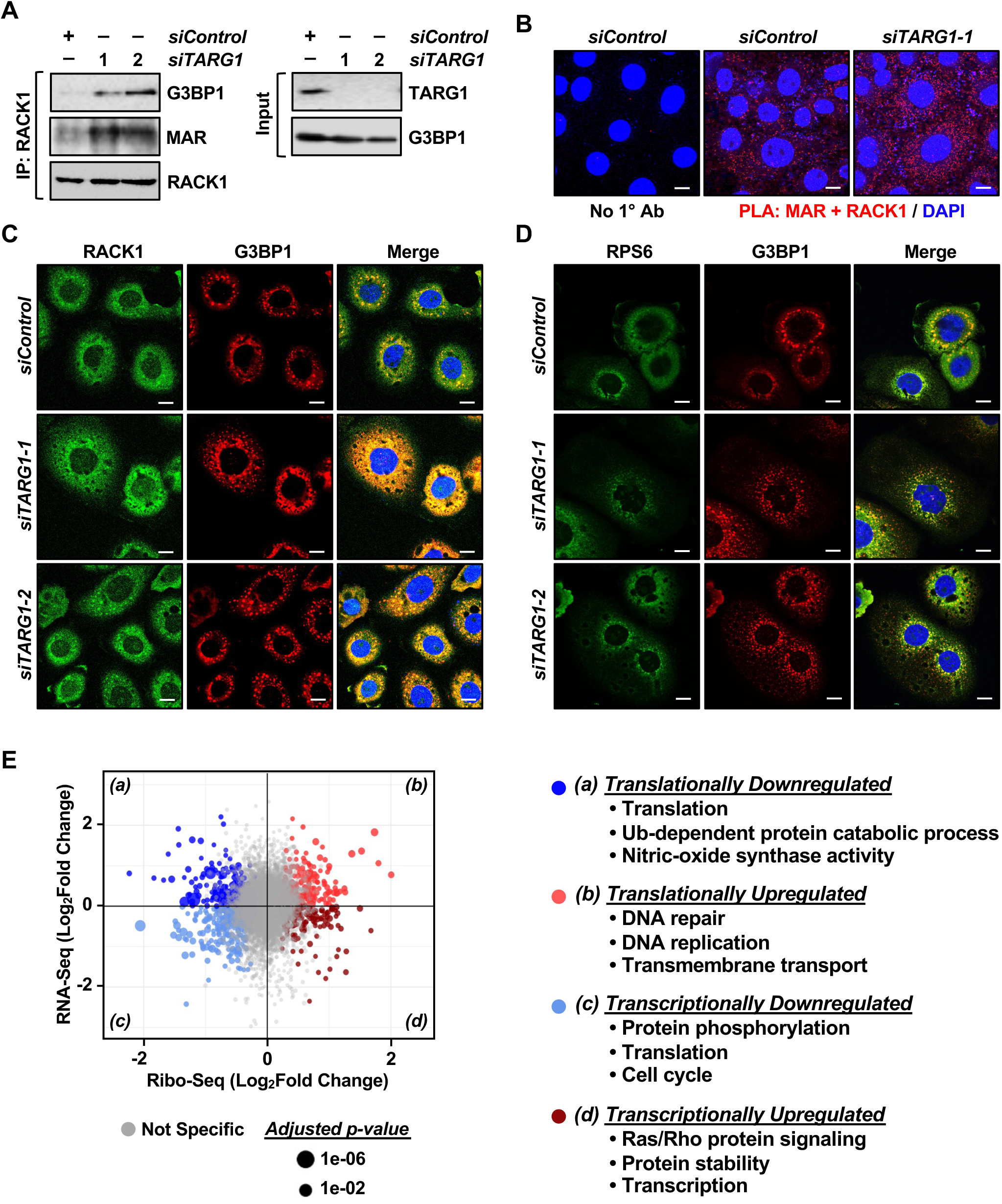
Depletion of TARG1 enhances stress granule assembly by increasing RACK1 MARylation. **(A and B)** siRNA-mediated *TARG1* depletion increases RACK1 MARylation and enhances RACK1 interaction with G3BP1 in OVCAR3 cells subjected to 15 minutes of treatment with 250 µM sodium arsenite (NaAsO_2_). (A) RACK1 was immunoprecipitated from OVCAR3 cells with siRNA-mediated knockdown of *TARG1* and subjected to immunoblotting for G3BP1, MAR, and RACK1. (B) PLA using MAR and RACK1 antibodies. DNA was stained with DAPI. Scale bar is 15 µm. **(C and D)** TARG1 knockdown increases the assembly of G3BP1-containing stress granules. Immunofluorescent staining assays of OVCAR3 cells with siRNA-mediated knockdown of *TARG1* subjected to 15 minutes of treatment with 250 µM sodium arsenite (NaAsO_2_). (C) RACK1 and G3BP1, (D) RPS6 and G3BP1. DNA was stained with DAPI. Scale bar is 15 µm. **(E)** Changes in mRNA translation upon depletion of TARG1. Scatter plot of fold changes in ribosome profiling and RNA-seq (OVCAR3 cells subjected to siRNA-mediated TARG1 knockdown vs. WT) comparing translational control and transcriptional control. Gene ontology enrichment analysis of the mRNAs regulated at transcriptional and translational levels are shown.

### Prolonged exposure to stressors reduces RACK1 MARylation and localization to stress granules

Our data support the conclusion that TARG1 deMARylates RACK1 to reduces stress granule assembly. Prior studies have indicated that TARG1 localizes to stress granules (Butepage et al., 2018; Zaja et al., 2020), so we investigated whether TARG1 deMARylates RACK1 within stress granules. Indeed, prolonged exposure to stressors that cause stress granule assembly, such as sodium arsenite (NaAsO_2_) or thapsigargin, reduced RACK1 MARylation (Fig. 6A). Moreover, PLAs using cells subjected to prolonged exposure (30 minutes) to sodium arsenite demonstrated that the increased stress granule assembly caused by this exposure increased the interaction of RACK1 with TARG1 (Fig. 6B, top; Suppl. Fig. S5H). Furthermore, PLAs also demonstrated that depletion of TARG1 blocks sodium arsenite-mediated deMARylation of RACK1 (Fig. 6B, bottom; Suppl. Fig. S5I). Finally, we performed PLAs to measure the interaction of RACK1 with TARG1 in OVCAR3 cells expressing GFP-tagged G3BP1 that we subjected to sodium arsenite treatment. We observed that TARG1-RACK1 complexes exists outside of stress granules near the GFP-G3BP1 puncta (Fig. 6C). Although the relationship between stress granule assembly and TARG1 interaction with RACK1 remains to be elucidated, these results support the role of deMARylation of RACK1 by TARG1 in the inhibition of stress granule assembly (Fig. 6D).

**Figure 6.**
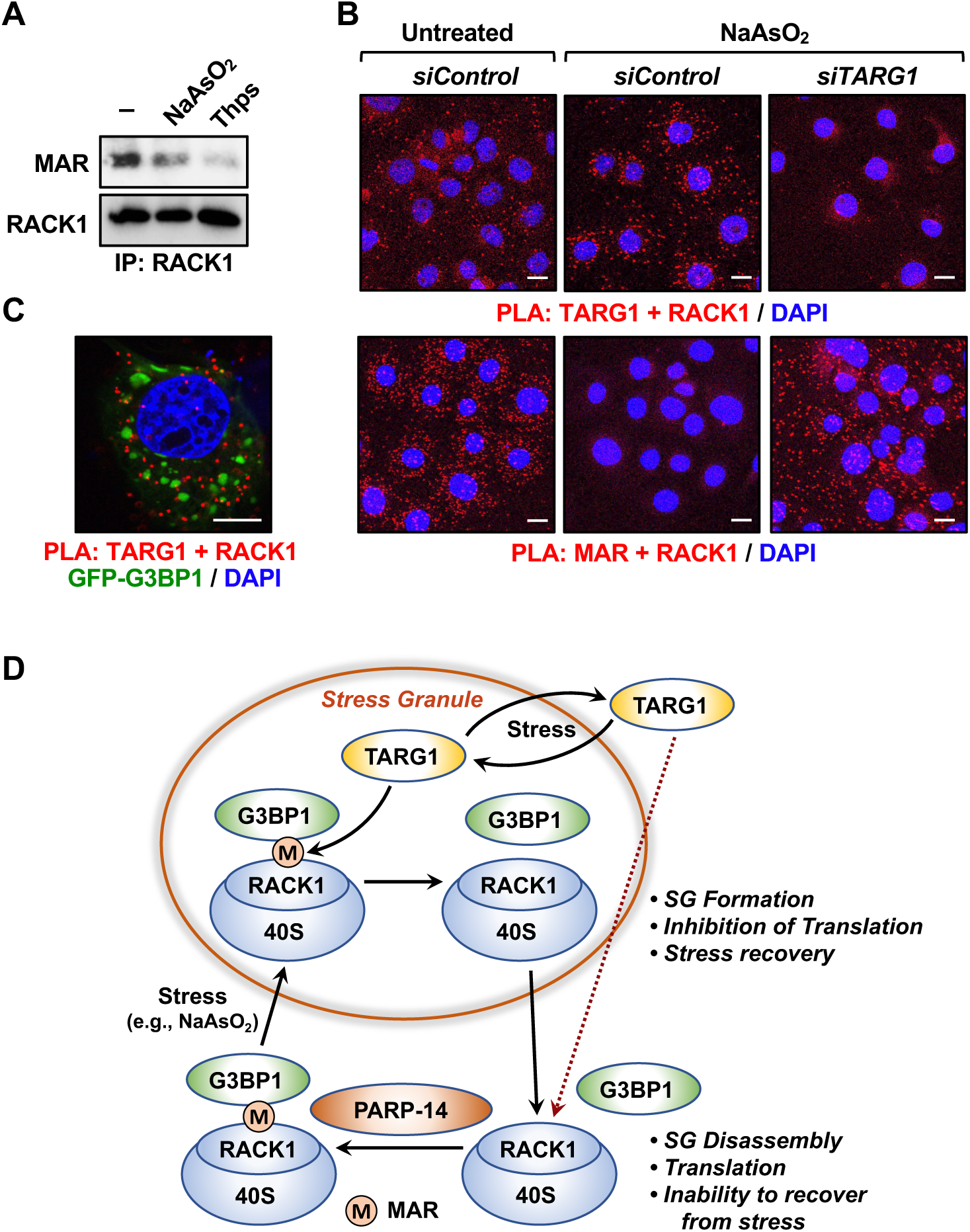
Prolonged exposure to stress reduces RACK1 MARylation. **(A and B)** Stress reduces RACK1 MARylation through TARG1. (A) RACK1 was immunoprecipitated from OVCAR3 cells treated with 250 µM sodium arsenite (NaAsO_2_) for 30 minutes or 250 nM thapsigargin (Thps) for 2 hours, and subjected to immunoblotting for MAR and RACK1. (B) PLA in OVCAR3 cells with siRNA-mediated knockdown of *TARG1* using TARG1 and RACK1 (*top*) or MAR and RACK1 (*bottom*) antibodies. The cells were treated with sodium arsenite (NaAsO_2_) treatment for 30 minutes. DNA was stained with DAPI. Scale bar is 15 µm. **(C)** Knockdown of *TARG1* increases the assembly of G3BP1-containing stress granules. PLA for TARG1 and RACK1 combined with immunofluorescent imaging of OVCAR3 cells expressing GFP-G3BP1 with siRNA-mediated knockdown of TARG1 and treated with 250 µM sodium arsenite (NaAsO_2_) for 15 minutes. DNA was stained with DAPI. Scale bar is 15 µm. Schematic of the mechanisms by which PARP14 and TARG1 regulate stress granule assembly through RACK1 MARylation. Additional details are provided in the text.

## Discussion

Recent studies have shown that ribosomal proteins and other components of the translation machinery are ADP-ribosylated (ADPRylated) in cancers (Challa et al., 2021a; Gibson et al., 2016; Zhen et al., 2017). Interestingly, MARylation of the translation machinery is a well-characterized outcome of intoxication by several human bacterial pathogens (e.g., *C. diphtheria* and *P. aeruginosa*, and *V. cholera*), whose toxins (diphtheria toxin, exotoxin A, and cholix toxin, respectively) MARylate host elongation factor-2 (eEF2), an essential component of the protein translation machinery, on a unique diphthamide residue in domain IV (Deng and Barbieri, 2008; Jorgensen et al., 2008). MARylation of eEF2 halts protein synthesis and causes cell death (Deng and Barbieri, 2008; Mateyak and Kinzy, 2013; Simon et al., 2014). Recent studies have begun to link ribosome biogenesis, ribosome function, and translation to cellular outcomes in cancers (Brighenti et al., 2015; Bustelo and Dosil, 2017; Dai and Lu, 2008; van Sluis and McStay, 2014). These published studies, in conjunction with the data that we present herein, suggest an intriguing link between ribosomal MARylation and ribosome function in cancer.

### PARP14 and TARG1 mediate site-specific MARylation and deMARylation of RACK1, respectively

We have previously shown that two specific sets of ribosomal proteins are MARylated: (1) “assembly factors” (e.g., RPS6, RPL24) located at the interface between the 60S and 40S ribosomal subunits and (2) “regulatory factors” (e.g., RACK1) located on the surface of the 40S subunit (Challa et al., 2021a). Moreover, we showed that PARP16, a tail-anchored endoplasmic reticulum-associated protein, is a MART that MARylates RPS6 and RPL24 in ovarian cancer cells to control the loading of mRNAs on ribosomes and their translation. Herein, we demonstrate that PARP14 is a MART that MARylates RACK1 in ovarian cancer cells to control stress granule formation and the regulation of translation under cellular stress conditions. In addition, we showed that TARG1, an Asp/Glu ADP-ribosyl hydrolase (Sharifi et al., 2013), deMARylates RACK1 to dissociate stress granules and return RACK1 and the 40S ribosomal subunit to the cytoplasm, allowing for a restoration of translation.

RACK1 is an integral component of the 40S ribosome (Rabl et al., 2011) and member of the tryptophan-aspartate repeat (WD-repeat) family of proteins (Adams et al., 2011). We mapped and confirmed three sites of MARylation at Asp 144, Glu 145, and Asp 203, which occur within blades 4 and 5 of β-propeller domain of RACK1. These residues are located in a highly accessible region of the ribosome (Fig. 1A), making them ideal candidates for regulation pf protein-protein interactions, such as those that might drive interactions with G3BP1 to bring the 40S ribosomal subunit into stress granules, as we observed.

### A RACK1 MARylation cycle drives stress granule dynamics in ovarian cancer cells

We observed that RACK1 is MARylated by PARP14 in ovarian cancer cells. Site- specific MARylation of RACK1 is required for stress granule formation and it promotes the colocalization of RACK1 to stress granules with key markers, such as G3BP1, eIF3η, and 40S ribosomal proteins. We also demonstrated that TARG1 reverses RACK1 MARylation and assembly of stress granules. We used ribosome profiling assays to identify changes in translation caused by dysregulated RACK1 MARylation. These results revealed that cells expressing RACK1-Mut, as well as cells with TARG1 depletion, exhibit decreased translation of mRNAs encoding proteins contributing to key pathways in ovarian cancer, including AKT1 and DNA repair. Collectively, our results support a PARP14/TARG1-regulated RACK1 MARylation cycle that controls stress granule assembly and disassembly in ovarian cancer cells (Fig. 6D). While previous studies have focused extensively on mechanisms of stress granule assembly, a unique aspect of our work is the discovery of a deMARylation-dependent mechanism that leads to the dissociation of stress granules.

How does site-specific MARylation of RACK1 allow it to engage and regulate the molecular pathway leading to stress granule formation? Although we have not addressed this question in detail, protein-linked ADPR moieties have been shown to alter the biochemical and structural properties of the proteins that contain them (Challa et al., 2021b; Gupte et al., 2017). Moreover, ADPR moieties can create binding sites for ADPR binding domains that drive protein-protein interactions (Challa et al., 2021b; Gupte et al., 2017). Both of these mechanisms may be relevant in the context of RACK1. Our results also demonstrate that disrupting polysome formation normalizes the differences in stress granule formation in RACK1-WT and RACK1-Mut expressing cells. These data, along with our observation that G3BP1 is less associated with polysomes in cells expressing RACK1-Mut, suggest that MARylation of RACK1 regulates stress granule assembly by affecting polysome function.

### PARP14 as a potential therapeutic target in ovarian cancer

Interest in drugging cytosolic MARTs for therapeutic purposes is growing (Peng et al., 2017; Yoneyama-Hirozane et al., 2017). Efforts are continuing to develop chemical inhibitors of MARTs, including inhibitors of PARP7 (Gozgit et al., 2021; Sanderson et al., 2023), PARP14 (Schenkel et al., 2021), and PARP16 (Bejan et al., 2022). In our studies, treatment with a PARP14 inhibitor (RBN012759) (Schenkel et al., 2021) phenocopied the effects of mutation of the sites of MARylation on RACK1, indicating that many of the effects of the PARP14 inhibitor were mediated through RACK1 MARylation. These results suggest that chemical inhibition of PARP14 may be a useful strategy for the treatment of ovarian cancer, which is supported by the results of our cell growth and xenograft tumor assays (Fig. 4, B and D; Suppl. Fig. S3, A, B, D, and G).

Collectively, the results that we presented here support the dynamic control of RACK1 MARylation during stress granule assembly, in response to stressors (Fig. 6D). In sum, we showed that PARP14-mediated MARylation of RACK1 controls stress granule assembly, while TARG1-mediated deMARylation of RACK1 reverses stress granule assembly. We also demonstrated that pharmacological inhibition of PARP14 sensitizes ovarian cancer cells to stress by inhibiting stress granule assembly. Our results define a new pathway in the control of stress granules in cancer and support the growing link between ribosomal MARylation and ribosome function in cancer.

## Supporting information

All Supplemental Figures 1-5

## Acknowledgements

The authors thank Subin Myong for technical assistance and Dr. Adi Gazdar for the HCC5044 cell line. The authors would also like to thank Dr. Anthony Leung and colleagues for sharing their related paper prior to submission.

## Author Contributions

Sridevi Challa – project conception, all main figure data generation and analyses, preparing figures, writing, editing, funding.

Tulip Nandu – computational analyses for integration of RNA-seq and Ribo-seq data, writing of methods.

Hyung Bum Kim – ADP-ribosyl hydrolase screen.

Xuan Gong – xenograft tumor assays.

Charles W. Renshaw – growth assays, imaging assays, and immunoblots in SKOV3 and HCC5044 cells.

Wan-Chen Li - Ribo-seq library generation.

Xinrui Tan – analysis of ribosome MARylation data and structural modeling.

Marwa W. Aljardali – PLAs and immunofluorescence assays.

Cristel V. Camacho - xenograft tumor assays, writing of methods.

Jin Chen – Ribo-seq library generation and data analysis, writing of methods.

1. W. Lee Kraus - project conception, data analyses, preparing figures, writing, editing, funding.

## Supplemental Figures

This manuscript contains a supplemental file with 5 supplemental figures, Figs. S1 through S5.

## Materials and Data Availability

### Materials availability

All cell lines and DNA constructs are available by request from W. Lee Kraus. The mono(ADP-ribose) detection reagent is available for purchase from EMD Millipore.

### Data and code availability

The ribosome profiling data sets generated specifically for this study can be accessed from the NCBI’s Gene Expression Omnibus (GEO) repository (http://www.ncbi.nlm.nih.gov/geo/) using the superseries accession number GSE245504. Computational scripts and pipelines are available from W.L.K and J.C., or on github (https://github.com/Kraus-Lab/RACK1_MARylation_Cycle).

## Materials and Methods

### Antibodies and chemicals

The custom recombinant antibody-like anti-MAR binding reagent (anti-MAR) was generated and purified in-house (now available from Millipore Sigma, MABE1076) (Gibson et al., 2017b). The other antibodies used were as follows: PARP14 (rabbit polyclonal; Sigma, HPA008846), PARP14 (mouse monoclonal; Santa Cruz, sc-377150), G3BP1 (Proteintech, 13057-2-AP), β-tubulin (Abcam, ab6046), RACK1 (Santa Cruz, sc-377150), RPS6 (Santa Cruz, sc-74459), eIF3η (Santa Cruz, sc-137214), puromycin (Millipore, MABE343), phospho-eIF2α (Cell Signaling, 9721), eIF2α (Cell Signaling, 9722), cleaved caspase-3 (Cell Signaling, 9661S), HA tag (mouse monoclonal; Sigma-Aldrich, H3663), HA tag (rabbit polyclonal; Abcam, ab9110), mouse IgG (Invitrogen, 10400C), goat anti-rabbit HRP-conjugated IgG (Pierce, 31460), and goat anti-mouse HRP-conjugated IgG (Pierce, 31430). The specialized chemicals used were thapsigargin (Tocris, 1138) and RBN012759 (MedChemExpress, HY-136979).

### Cell culture

OVCAR3, SKOV3, and HEK-293T cells were purchased from the American Type Cell Culture (ATCC). HCC5044 cells were obtained from Dr. Adi Gazdar (Thu et al., 2017). OVCAR3 and SKOV3 were maintained in RPMI (Sigma-Aldrich, R8758) supplemented with 10% fetal bovine serum and 1% GlutaMax (Thermo Fisher Scientific, 35050061). HCC5044 and HEK-293T cells were cultured in DMEM (Sigma-Aldrich, D5796) supplemented with 10% fetal bovine serum. Fresh cell stocks were regularly replenished from the original stocks every few months, verified for cell type identity using the GenePrint 24 system (Promega, B1870), and confirmed as mycoplasma-free every three months using a commercial testing kit.

### Cell treatments

Ovarian cancer cells were treated with the various chemicals and inhibitors as described herein. To inhibit PARP14, the cells were treated with RBN012759 (10 μM; MedChemExpress, HY-136979) for 24 hours. For induction of stress granules, the cells were treated with sodium arsenite (NaAsO_2_; 250 μM; Sigma, S7400) or thapsigargin (250 nM; Tocris, 1138) for the times indicated in the figure legends. Both are commonly used experimentally to induce stress granule formation. NaAsO_2_ causes oxidative stress and protein misfolding, leading to eIF2α phosphorylation, stalled translation, and stress granule assembly (Bernstam and Nriagu, 2000). In contrast, thapsigargin depletes Ca^2+^ stores in the endoplasmic reticulum (ER), thereby causing ER stress and stress granule assembly (Thastrup et al., 1990). We routinely use both agents in our experiments examining stress granule formation.

### Generation of vectors for inducible knockdown or ectopic expression

We purchased vectors for shRNA-mediated knockdown of mRNAs and generated vectors for ectopic expression of proteins using the oligonucleotide primers described below. All constructs were verified by sequencing.

#### Vectors expressing shRNAs targeting RACK1

The pTRIPZ vector for expressing an shRNA targeting human *RACK1* was purchased from Horizon Discovery (RHS4696-200769910) along with the control pTRIPZ vector, which were used as described previously (Ryu et al., 2018).

#### Vectors for ectopically expressing RACK1

A plasmid for Dox-inducible expression of C-terminal HA epitope-tagged RACK1 was generated using a cDNA for *RACK1* that was amplified from pCMV3-RACK1 (Sino Biologicals, HG16196-CY) and subcloned into the pInducer20 vector (Addgene, plasmid no. 44012). The mutations corresponding to the MARylation sites of RACK1 (D144N/E145Q/D203N) were introduced into the pCMV3- RACK1 plasmid using the QuikChange Site-Directed Mutagenesis kit (Agilent).

### Oligonucleotides used for cloning

#### Cloning primers for pInducer-RACK1

- Cloning forward: 5’-TCCGCGGCCCCGAACTAGTGATGACTGAGCAGATGACC-3’

- Cloning reverse: 5’-GTTTAATTAATCATTACTACTTAGGCGTAGTCAGGCAC-3’

#### Primers for generating RACK1-Mut

- D144N forward: 5’-TACACTGTCCAGAATGAGAGCCACTCAGAG-3’

- D144N reverse: 5’-CTCTGAGTGGCTCTCATTCTGGACAGTGTA-3’

- E145Q forward: 5’-TACACTGTCCAGGATCAGAGCCACTCAGAG-3’

- E145Q reverse: 5’-CTCTGAGTGGCTCTGATCCTGGACAGTGTA-3’

- D203N forward: 5’-GACTGTCTCTCCAAATGGATCCCTCTGTG-3’

- D203N reverse: 5’-CACAGAGGGATCCATTTGGAGAGACAGTC-3’

### Generation of cell lines with inducible knockdown or ectopic expression

Cells were infected with lentiviruses for inducible knockdown or ectopic expression. We generated lentiviruses by transfection of the pTRIPZ or pInducer20 constructs described above, together with an expression vector for the VSV-G envelope protein (pCMV-VSV-G, Addgene plasmid no. 8454), an expression vector for GAG-Pol-Rev (psPAX2, Addgene plasmid no. 12260), and a vector to aid with translation initiation (pAdVAntage, Promega) into HEK-293T cells using GeneJuice transfection reagent (Novagen, 70967) according to the manufacturer’s protocol. The resulting viruses were used to infect the ovarian cancer cells in the presence of 7.5 μg/mL polybrene for 24 hours or 48 hours after the initial transfection. Stably transduced cells were selected with puromycin (Sigma, P9620; 1 μg/mL) for pTRIPZ or G418 sulfate (Sigma, A1720; 500 µg/mL) for pInducer20. For inducible knockdown or expression, the cells were treated with 1 µg/mL doxycycline (Dox) for 48 hours.

### siRNA-mediated knockdown

For siRNA-mediated knockdown, the siRNAs listed below were transfected into ovarian cancer cells at a final concentration of 30 nM using Lipofectamine RNAiMAX reagent (Invitrogen, 13778150) according to the manufacturer’s instructions. The cells were used for various assays 48 hours after siRNA transfection. The siRNAs for knocking down MARTs were described previously (Challa et al., 2021a). The control siRNA and the siRNAs for knocking down hydrolases were purchased from Sigma as follows:

- GDAP2: siRNA1: SASI_Hs01_00119609; siRNA2: SASI_Hs01_00119610

- MacroD1: siRNA1: SASI_Hs01_00236121; siRNA2: SASI_Hs01_00236122

- MacroD2: siRNA1: SASI_Hs01_00140117; siRNA2: SASI_Hs01_00140118

- NUDT16: siRNA1: SASI_Hs01_00032889; siRNA2: SASI_Hs01_00032890

- TARG1: siRNA1: SASI_Hs01_00165859; siRNA2: SASI_Hs02_00364965

- PARP14: siRNA1: SASI_Hs02_00350199; siRNA2: SASI_Hs01_00178227

### Preparation of whole cell lysates

Cells were cultured and treated as described above before the preparation of cell extracts. At the conclusion of the treatments, the cells were washed twice with ice-cold PBS and resuspended in Lysis Buffer (20 mM Tris-HCl pH 7.5, 150 mM NaCl, 1 mM EDTA, 1 mM EGTA, 1% NP-40, 1% sodium deoxycholate, 0.1% SDS) containing 1 mM DTT, 250 nM ADP-HPD, 10 μM PJ-34, 1x complete protease inhibitor cocktail (Roche, 11697498001) and phosphatase inhibitors (10 mM sodium fluoride, 2 mM sodium orthovanadate, and 10 mM β- glycerophosphate). The cells were vortexed for 30 seconds in Lysis Buffer and then centrifuged at full speed for 15 minutes at 4°C in a microcentrifuge to remove the cell debris.

### Isolation of polysomes

To isolate polysomes, 5 million cells were plated in 15 cm diameter dishes and treated as described above 24 hours prior to the assay. Polysomes were isolated from the cells using a previously described protocol (Morita, 2013) with some modifications. Briefly, the cells were treated with 100 µg/mL cycloheximide for 10 minutes, then washed three times with ice-cold PBS containing 100 µg/mL cycloheximide. The cells were collected by gentle scraping in 500 µL Polysome Lysis Buffer (15 mM Tris HCl pH 7.4, 15 mM MgCl_2_, 250 mM NaCl, 1% Triton X-100 in DEPC water) supplemented with 1 mM DTT, 100 µg/mL cycloheximide, and 400 U/mL RNase inhibitor (Promega; N2611), as well as the protease, phosphatase, PARG (ADP-HPD), and PARP (PJ-34) inhibitors noted above. The resuspended cells were vortexed for 30 seconds and centrifuged at full speed for 15 minutes at 4°C in a microcentrifuge. Five percent of the lysate was aliquoted to be used as input for measuring the steady state mRNA or protein levels. RNA content was measured by reading the absorbance at 260 nm, and equal amounts of RNA were loaded onto 15-50% sucrose gradients. The gradients were centrifuged at 125,000 x g for 2 hours at 4°C in a Beckman coulter Optima L-80 XP ultracentrifuge using a SW60Ti rotor. The gradient was collected as 250 µL fractions in 2 mL microfuge tubes. The RNA content in these fractions were measured by reading the absorbance at 260 nm and the peaks corresponding to monosomes and polysomes were noted.

For SDS-PAGE analyses, the proteins were precipitated from the fractions using methanol-chloroform. Briefly, 900 µL of methanol were added to each 250 µL fraction with mixing by inversion, then 225 µL of chloroform were added with mixing by vortexing. Finally, 675 µL of ddH_2_O were added to the tubes, followed by vortexing until a precipitate was observed. The samples were centrifuged at full speed for 5 minutes at 4°C in a microcentrifuge. The upper phase was removed by aspiration and the protein pellet was washed by adding 750 µL methanol with gentle mixing. The protein pellet was re-collected by centrifugation at full speed for 5 minutes at 4°C in a microcentrifuge. After the protein pellets were allowed to air dry briefly, they were dissolved in 1x SDS-PAGE loading solution, heated at 50°C for 10 minutes, and heated to 95°C for SDS-PAGE and subsequent immunoblotting.

### Immunoblotting

The protein concentrations of the cell lysates were determined using a Bio-Rad Protein Assay Dye Reagent (Bio-Rad, 5000006). Volumes of lysates containing the same amount of total protein were heated to 95°C for 5 minutes after addition of 1/4 volume of 4x SDS-PAGE Loading Solution (250 mM Tris, pH 6.8, 40% glycerol, 0.04% Bromophenol Blue, 4% SDS). The lysates or the polysome fractions described above, were run on polyacrylamide-SDS gels, and transferred to nitrocellulose membranes. After blocking with 5% nonfat milk in TBST, the membranes were incubated with the primary antibodies described above in TBST with 0.02% sodium azide, followed by anti-rabbit HRP-conjugated IgG (1:5000) or anti-mouse HRP-conjugated IgG (1:5000). Immunoblot signals were detected using an ECL detection reagent (Thermo Fisher Scientific, 34577, 34095).

### Puromycin incorporation assays

Protein synthesis was determined using puromycin incorporation assays as previously described (Schmidt et al., 2009). Briefly, OVCAR3 cells with Dox-inducible knockdown were plated at 50% confluence in 6-well plates. Forty-eight hours later, the cells were treated with 10 μg/mL puromycin for 15 minutes at 37°C. Whole cell lysates were prepared from these cells as described above and puromycin incorporation was visualized by immunoblotting using an antibody against puromycin.

### RNA isolation and reverse transcription-quantitative real-time PCR (RT-qPCR)

OVCAR3 cells were transfected with different siRNAs as described above and total RNA was isolated using the Qiagen RNeasy Plus Mini kit (Qiagen, 74136) according to the manufacturer’s protocol. Total RNA was reverse transcribed using oligo(dT) primers and MMLV reverse transcriptase (Promega, PR-M1705) to generate cDNA. The cDNA samples were subjected to RT-qPCR using gene-specific primers listed below. Target gene expression was normalized to the expression of *RPL19* mRNA. All experiments were performed a minimum of three times with independent biological replicates to ensure reproducibility and a statistical significance of at least p < 0.05. Statistical differences between control and experimental samples were determined using the Student’s t-test.

### RT-qPCR primers

- RPL19 forward: 5’-ACATCCACAAGCTGAAGGCA-3’

- RPL19 reverse: 5’-TGCGTGCTTCCTTGGTCTTA-3’

- GDAP2 forward: 5’-AGTTCTGGAATGATGACGACTCG-3’

- GDAP2 reverse: 5’-GTGGGTGTCGATACAGGTCAG-3’

- MACROD1 forward: 5’-CCAAAACCAGTTTCTTTGGGAG-3’

- MACROD1 reverse: 5’-CAGATTCCATCTACCACATCC-3’

- MACROD2 forward: 5’-TGTGCTAGTTACTACAGAGCCA-3’

- MACROD2 reverse: 5’-CCCCATCATAGTTCACCTGCC-3’

- NUDT16 forward: 5’-TACGGGAAGGGCGTGTATTTC-3’

- NUDT16 reverse: 5’-GCCACGAACACCGCCTTAT-3’

- TARG1 forward: 5’-ATCTGCCAGCAGAACTTTGA-3’

- TARG1 reverse: 5’-AACATCGTGTGGGTCTGCGTGT-3’

### Immunofluorescent staining and confocal microscopy of cultured cells

The following microscopy-based protocols for cultured cells were used to assess stress granule formation, protein localization, and protein MARylation in cells.

#### Immunofluorescent staining

OVCAR3 cells were seeded on 8-well chambered slides (Thermo Fisher, 154534) one day prior to the experiment. The cells were washed once with PBS, fixed in 4% paraformaldehyde for 15 minutes at room temperature, and washed three times with PBS. The cells were permeabilized for 10 minutes at -20°C using ice cold methanol, washed three times with PBS, and incubated for 1 hour at room temperature in Blocking Solution (PBS containing 1% BSA, 10% FBS, 0.3M glycine and 0.1% Tween-20). The fixed cells were incubated with a mixture of the primary antibodies in PBS overnight at 4°C, followed by three washes with PBS. The cells were then incubated with a mixture of Alexa Fluor 594 donkey anti-rabbit IgG (ThermoFisher, A-21207) and Alexa Fluor 488 goat anti-mouse IgG (ThermoFisher, A-11001) each at a 1:500 dilution in PBS for 1 hour at room temperature. After incubation, the cells were washed three times with PBS. Finally, coverslips were placed on cells coated with VectaShield Antifade Mounting Medium with DAPI (Vector Laboratories, H-1200) and images were acquired using an inverted Zeiss LSM 880 confocal microscope.

#### Proximity ligation assays (PLAs)

Proximity ligation assays were performed using a Duolink proximity ligation kit (Sigma-Aldrich, DUO92008) following the manufacturer’s instructions. Briefly, cells were plated on sterilized microscope cover glass (Fisherbrand, 12CIR-1.5). At the end of treatments, the cells were washed once with PBS, fixed in 4% paraformaldehyde for 15 minutes at room temperature, and washed three times with PBS. The cells were permeabilized for 10 minutes at -20°C using ice cold methanol, washed three times with PBS, and incubated in Duolink Blocking Solution for 1 hour at 37°C in a humidified chamber. Excess Blocking Solution was removed by tapping and the cells were incubated in the primary antibody pairs: mouse monoclonal RACK1 (1:200) or HA (1:200) antibodies and rabbit polyclonal MAR (1:200) or G3BP1 (1:1000) or TARG1 (1:500) antibodies in Duolink Antibody Diluent. The slides were incubated overnight at 4°C in a humidified chamber.

Following the overnight incubation, the cells were washed twice for 5 minutes each with Wash Buffer A (10 mM Tris-HCl pH 7.4, 150 mM NaCl, and 0.05% Tween). The slides were then incubated with Ligation Solution (1:40 dilution of the ligase in 1x Ligation Buffer) for 30 minutes at 37°C in a humidified chamber, followed by two washes with Wash Buffer A for 5 minutes each. The cells were then incubated in the Amplification Solution (1:80 dilution of the Polymerase in 1x Amplification Buffer) for 100 minutes at 37°C in a humidified chamber protected from light. The cells were washed twice with Wash Buffer B (200 mM Tris-HCl pH 7.5, 100 mM NaCl) for 10 minutes each followed by a wash with 0.01x Wash Buffer B for 2 minutes. The stained cells were mounted on microslides using VectaShield Antifade Mounting Medium (Vector laboratories, H-1200-10) with DAPI DNA stain. Images were acquired using an inverted Zeiss LSM 880 confocal microscope.

#### Image analysis

The fluorescence intensities captured by the confocal imaging were analyzed by Fiji ImageJ software (Schindelin et al., 2012). The intensity and contrast of the images were further adjusted in Microsoft Powerpoint and same changes were applied to all of the samples in each condition. The number of G3BP1 foci (i.e., stress granules) or PLA foci were normalized to the number of nuclei (i.e., the number of cells) to determine the average number of foci/cell.

### siRNA screen to identify MARTs that mediate RACK1 MARylation

OVCAR3 cells were plated into 24-well plates containing the microscope cover glass and transfected with 30 nM each of the PARP mRNA-targeting siRNAs as described above. Two different siRNAs per PARP mRNA were used. Forty-eight hours after transfection, RACK1 MARylation levels were determined using proximity ligation assays as described above.

### Determination of PARP14 autoMARylation

Cells were grown in 15 cm plates were treated with RBN012759 for 24 hours. Twenty-four hours after transfection, the cells were harvested in ice-cold PBS and then lysed in 500 mM Lysis Buffer (50 mM Tris-HCl pH 7.5, 500 mM NaCl, 1 mM EDTA, 1% IGEPAL CA-630, 10% glycerol, and 1 mM DTT) containing 1x complete protease inhibitor cocktail, phosphatase inhibitors, PARG inhibitor, and PARP inhibitor as described above. Volumes of lysate containing equal amounts of total protein were used to immunoprecipitate PARP14 by incubating with 2 µg of a mouse monoclonal antibody against PARP14 (Santa Cruz, sc-377150), and protein G agarose beads overnight at 4°C with gentle mixing on a nutator at 4°C. After incubation overnight, the beads were washed three times for 5 minutes each at 4°C with 500 mM Lysis Buffer. The beads were heated to 95°C in 1x SDS-PAGE loading buffer. The samples were run on an 8% SDS-PAGE gel and transferred to a nitrocellulose membrane for immunoblotting as described above. Autoactivation of PARP14 was determined by immunoblotting with a MAR detection reagent (Millipore Sigma, MABE1076).

### Determination of RACK1 MARylation

Cells were transfected with pCMV3-RACK1 for expressing HA-tagged wild-type (WT) or MARylation site mutant (Mut) RACK1 using GeneJuice transfection reagent. Forty-eight hours after transfection, the cells were harvested in ice-cold PBS and then lysed in 500 mM Lysis Buffer (50 mM Tris-HCl pH 7.5, 500 mM NaCl, 1 mM EDTA, 1% IGEPAL CA-630, 10% glycerol, and 1 mM DTT) containing 1x complete protease inhibitor cocktail, phosphatase inhibitors, PARG inhibitor, and PARP inhibitor as described above. Volumes of lysate containing equal amounts of total protein were used to immunoprecipitate RACK1 by incubating with mouse monoclonal antibody against HA and protein G agarose beads (Thermo Fisher, 15920010) at 4°C with gentle mixing. After incubation overnight, the beads were washed three times for 5 minutes each at 4°C with 500 mM Lysis Buffer. The beads were heated to 95°C in 1x SDS-PAGE loading buffer. The samples were run on a 12% SDS-PAGE gel and transferred to a nitrocellulose membrane for immunoblotting with a MAR detection reagent (Millipore Sigma, MABE1076).

### Co-immunoprecipitation of RACK1 with G3BP1

The cells were cultured in 15 cm diameter dishes and subjected to treatments as described above. They were washed twice with ice-cold PBS and then lysed in IP Lysis Buffer (50 mM Tris-HCl pH 7.5, 150 mM M NaCl, 1.0 mM EDTA, 1% NP-40 and 10% glycerol, supplemented with fresh 1 mM DTT, 250 nM ADP-HPD, 10 μM PJ-34, 1x complete protease inhibitor cocktail (Roche, 11697498001) and phosphatase inhibitors (10 mM sodium fluoride, 2 mM sodium orthovanadate, and 10 mM β-glycerophosphate). Volumes of lysate containing equal amounts of total protein were used to immunoprecipitate RACK1 by incubating with 2 µg of anti-HA antibody (ectopically expressed RACK1) or anti-RACK1 antibody (endogenous RACK1) and protein G beads with gentle mixing overnight on a nutator at 4°C. After incubation, the beads were washed five times for 5 minutes each at 4°C with IP Lysis Buffer. The beads were then heated to 95°C for 5 minutes in 1x SDS-PAGE loading buffer, and the immunoprecipitated proteins were run on a 10% PAGE-SDS gel, transferred to a nitrocellulose membrane, and immunoblotted as described above.

### Co-immunoprecipitation of G3BP1 interacting proteins

The cells were cultured in 15 cm diameter dishes and subjected to treatments as described above. They were washed twice with ice-cold PBS and then collected in Low Salt IP Lysis Buffer (50 mM Tris-HCl pH 7.5, 135 mM NaCl, 1.0 mM EDTA, 1% NP-40 and 10% glycerol, supplemented with fresh 1 mM DTT, 250 nM ADP-HPD, 10 μM PJ34, 1x complete protease inhibitor cocktail). The cells were vortexed for 30 seconds and cell debris was cleared by centrifugation for 10 minutes at 4°C at full speed in a microcentrifuge. The protein concentrations in the supernatants were measured using a Bradford assay. And an equal amount of total protein was used for each IP condition. The cell lysates were incubated with 2 µg of a rabbit polyclonal G3BP1 antibody and protein A agarose beads overnight at 4°C with gentle mixing. The beads were then heated to 95°C for 5 minutes in 1x SDS-PAGE loading buffer, and the immunoprecipitated proteins were run on a 12% PAGE-SDS gel, transferred to a nitrocellulose membrane, and immunoblotted as described above using the indicated antibodies.

### Ribosome profiling (Ribo-seq) and RNA-sequencing (RNA-seq) library preparation and sequencing

#### Ribo-seq library generation

For performing ribosome profiling (Ribo-seq), ribosome-protected footprints were prepared for sequencing as described in a recently updated protocol of ribosome profiling (Chen et al., 2020; McGlincy and Ingolia, 2017). Briefly, the cells were plated, grown, and subjected to the various treatments and experimental conditions indicated as described above. They were the rapidly harvested and lysed. Clarified cell lysates were treated with RNase I (Invitrogen) to digest RNA not protected by ribosomes. The 80S ribosomes were isolated by centrifuging lysates through a 34% sucrose cushion at 100,000 x *g* for 1 hour at 4°C. The RNA was purified from the ribosome pellet using the Direct-zol RNA kit (Zymo Research). It was then resolved by electrophoresis through a denaturing gel, and the fragments corresponding to 28 to 34 bp were extracted from the gel.

The 3’ ends of the ribosome footprint RNA fragments were treated with T4 polynucleotide kinase (New England Biolabs, M0201) to allow ligation of a pre-adenylated DNA linker with T4 Rnl2(tr) K227Q (New England Biolabs, M0351S). The DNA linker used incorporates sample barcodes to enable library multiplexing, as well as unique molecular identifiers (UMIs) to enable removal of duplicated sequences. To separate ligated RNA fragments from unligated DNA linkers, 5’-deadenylase (Epicentre, DA11101K) was used to deadenylate the pre-adenylated linkers, which were then degraded by the 5’-3’ ssDNA exonuclease RecJ (NEB, M0264S). After rRNA reduction using the riboPOOL rRNA depletion kit (siTOOLs Biotech, *H. sapiens* pool), the RNA-DNA hybrid was used as a template for reverse transcription, followed by circularization with CircLigase (Epicentre, CL4111K). Finally, PCR of the cDNA circles was used to attach suitable adapters and indices for Illumina sequencing.

#### RNA-seq library generation

RNA-seq libraries were generated from total RNA obtained from the input samples from the Ribo-seq experiments described above (i.e., lysates without RNase digestion) using the TrueSeq Stranded Total RNA Library Prep (Illumina, 20020596).

#### Ribo-seq and RNA-seq library sequencing

The Ribo-seq and RNA-seq libraries were subjected to QC analyses using an Agilent TapeStation and sequenced using an Illumina HiSeq 2000.

### Ribo-seq and RNA-seq data analysis

#### Ribo-seq data analysis

For the ribosome profiling analysis, we used hg19/GRCh37 for the genome assembly and Gencode v.24 for the transcriptome reference. For processing of ribosome profiling data, linker sequences were removed from sequencing reads and the samples were de-multiplexed using FASTX-clipper and FASTX-barcode splitter (FASTX-Toolkit). Unique molecular identifiers and sample barcodes were then removed from reads using a custom Python script (available form J.C. or W.L.K upon request). Reads aligning to rRNAs and contaminants were filtered out using Bowtie v1.1.2, and all remaining reads were aligned to the custom transcriptome described above with Tophat v.2.1.1 (Kim et al., 2013) using --b2-very-sensitive --transcriptome-only --no-novel-juncs --max-multihits=64 flags. These alignments were assigned a specific P-site nucleotide using a 12-nt offset from the 3’ end of reads. Read counting and gene expression (RPKM) calculations are performed in Python 2.7 using Plastid (Dunn and Weissman, 2016) and differential expression analysis was done by DESeq2 (Love et al., 2014).

#### RNA-seq data analysis

The raw data were subjected to QC analyses using the FastQC tool (Andrews, 2010). We used the hg19/GRCh37 genome assembly and Gencode v.24 for the transcriptome reference to analyze RNA-seq data. For RNA-seq analysis, the reads aligned to rRNAs and contaminants were filtered out using Bowtie v1.1.2 (Langmead et al., 2009) and all remaining reads were aligned to the custom transcriptome with Tophat v.2.1.1 (Kim et al., 2013) using --b2-very-sensitive --transcriptome-only --no-novel-juncs --max-multihits=64 flags. Read counting and gene expression (RPKM) calculations are performed in Python 2.7 using Plastid (Dunn and Weissman, 2016) and differential expression analysis was done by DESeq2 (Love et al., 2014).

### Integration of Ribo-seq and RNA-seq data

#### Regulation of mRNA translation by RACK1 MARylation

Translation efficiency was calculated as ribosome profiling RPKM/RNA-seq RPKM. Heat maps were generated using Java TreeView (Saldanha, 2004) for the genes with significantly different translational efficiency in the RACK1-mutant. Gene ontology analyses were determined using the Database for Annotation, Visualization and Integrated Discovery (DAVID) Bioinformatics Resources website for gene ontology analysis (Huang da et al., 2009) for genes with significantly different translational efficiency in the RACK1-mutant.

#### Changes in mRNA translation upon depletion of TARG1

Scatter plot of fold changes in ribosome profiling and RNA-seq (i.e., OVCAR3 cells subjected to siRNA-mediated *TARG1* knockdown vs. control knockdown) comparing translational control and transcriptional control was generated using custom R script. Gene ontology enrichment analysis of the genes regulated at the transcriptional and translational levels (FC > 2) in each quadrant were generated using DAVID (Huang da et al., 2009)

### Cell growth assays

#### Cell growth assays for OVCAR3 cells with ectopic expression of RACK1

OVCAR3 cells with Dox-inducible knockdown and re-expression of RACK1 were plated at a density of 2,000 cells per well in a 96-well plate in growth medium containing 0.5 μg/mL puromycin, 200 μg/mL G418, and 1 μg/mL Dox. Twenty-four hours after plating the cells, the growth medium was replaced with fresh medium supplemented with 0.5 μg/mL puromycin, 200 μg/mL G418, and 1 μg/mL Dox in the presence of vehicle or 3 nM thapsigargin. The cells were grown for the indicated amount of time. At the end of the indicated times, cells were fixed with 4% paraformaldehyde for 15 minutes, washed with water, and stored at 4°C. The fixed cells were then stained with crystal violet (0.5% crystal violet in 20% methanol) for 30 minutes with gentle agitation at room temperature. The stained cells were washed with water and air dried. The crystal violet was then dissolved in 10% acetic acid and the absorbance at 590 nm was measured using a spectrophotometer. The absorbance of a blank well was subtracted from the samples and the values were normalized to the values at Day 0. Three independent biological replicates were performed to ensure reproducibility. Statistical differences were determined using two-way ANOVA.

#### Cell growth assays for combined PARP14 inhibitor and thapsigargin or carboplatin treatment

Ovarian cancer cells were plated at a density of 2,000 cells per well in 96-well plates. Twenty-four hours later, the cells were treated with single or combined treatments of 10 μM PARP14 inhibitor and 3 nM thapsigargin or 5 µM carboplatin for the indicated amount of time. At the end of the indicated times, the cells were processed for the crystal violet staining assay as described above.

### Xenograft experiments in mice

All animal experiments were performed in compliance with the Institutional Animal Care and Use Committee (IACUC) at the UT Southwestern Medical Center. Female NOD scid gamma (NSG) mice at 6-8 weeks of age were used. To establish ovarian cancer xenografts, 5 to10 x 10^6^ of OVCAR3 parental cells, or OVCAR3 cells engineered for Dox-inducible expression of RACK1 (WT or Mut) were injected subcutaneously in 100 μL into the flanks of mice in a 1:1 ratio of PBS and Matrigel (Fisher, CB 40230). All tumors were monitored until they reached an average volume of 100 mm^3^ to initiate experiment. For the experiment with cells ectopically expressing RACK1, mice were placed on a Dox containing diet (625 mg/kg; Envigo). For the PARP14 inhibitor experiments, mice were randomized into vehicle or PARP14 inhibitor treatment groups. Mice were treated with PARP14 inhibitor (MedChemExpress, HY- 136979) at a dose of 50 mg/kg diluted in 4% DMSO, 5% PEG 300, 5% Tween-80 in PBS or an equal volume of vehicle intraperitoneally (i.p.) daily for 5 days on, then 2 days off.

The weight of the mice was monitored once per week and tumor growth measured using electronic calipers approximately once a week. The tumor volumes were calculated using a modified ellipsoid formula: tumor volume = ½(length x width^2^). The xenograft experiments were carried out until the mice reached the end-point for euthanasia as required by IACUC. At the end of the experiment, the mice were euthanized to collect the xenograft tissue. The tissue was cut into several small pieces and separate portions were either snap-frozen in liquid nitrogen or fixed using 4% paraformaldehyde. The frozen tissues were pulverized using a tissue mill and lysed in Whole Cell Lysis Buffer (20 mM Tris-HCl pH 7.5, 150 mM NaCl, 1 mM EDTA, 1 mM EGTA, 1% NP-40, 1% sodium deoxycholate, 0.1% SDS, 1 mM DTT, 250 nM ADP-HPD, 10 mM PJ-34 supplemented with protease and phosphatase inhibitors) for extraction of protein. The protein samples were analyzed by immunoblotting as described above.

### Quantification and statistical analyses

All sequencing-based genomic experiments were performed a minimum of two times with independent biological samples. Statistical analyses for the genomic experiments were performed using standard genomic statistical tests as described above. All gene specific qPCR-based experiments were performed a minimum of three times with independent biological samples. Statistical analyses were performed using GraphPad Prism 9. All tests and p-values are provided in the corresponding figures or figure legends.

