## Supplementary material for "A PARP14/TARG1-Regulated RACK1 MARylation Cycle Drives Stress Granule Dynamics in Ovarian Cancer Cells": All Supplemental Figures 1-5

**Challa et al. (2024)**

**CONTENTS:**

• Supplemental Figures S1 through S5
• Supplemental Figure Legends S1 through S5

*<Fig. S1 is on the next page>*

**Supplemental Figure S1. RACK1-Mut inhibits the formation of G3BP1 foci.**
**(A and B)** RACK1 is MARylated. Endogenous RACK1 was immunoprecipitated from OVCAR3 cells transfected with siRNAs targeting a control sequence or *RACK1* and subjected to immunoblotting for MAR and RACK1. Each bar in (B) represents the mean + SEM of the levels of MARylated RACK1 and RACK1 in the immununoprecipitates of RACK1 (n = 3, Student's t-test, * $p < 0.05$ and ** $p<0.01$).
**(C)** Quantification of multiple experiments like the one shown in Figure 2A. Each bar represents the mean + SEM of the relative abundance of G3BP1 in HA - RACK1 immunoprecipitates (n = 3, Student's t-test, *** $p<0.001$).
**(D)** Quantification of multiple experiments like the one shown in Figure 2B. Each bar represents the mean + SEM of G3BP1-HA (RACK1) PLA foci from two biological replicates (n = 3, Student's t-test, ** $p < 0.01$).
**(E)** Quantification of multiple experiments like the one shown in Figure 2E. Each bar represents the mean + SEM of the relative abundance of eIF3η and RPS6 in G3BP1 immunoprecipitates (n = 3, Student's t-test, ** $p<0.01$).
**(F)** Quantification of multiple experiments like the one shown in Figure 2F. Each bar represents the mean + SEM of distinct G3BP1 foci from three biological replicates (n = 3, Student's t-test, ** $p < 0.01$).
**(G and H)** Loss of RACK1 MARylation inhibits G3BP1 localization to stress granules and its interaction with translation factors that are key components of stress granules. Immunofluorescent staining assays of OVCAR3 cells with Dox-induced knockdown of endogenous RACK1 and re-expression of exogenous RACK1 subjected to 15 minutes of treatment with 250 μM sodium arsenite ($NaAsO_2$). Staining for (A) RPS6 and G3BP1, (B) eIF3η and G3BP1. DNA was stained with DAPI. Scale bar is 15 μm.
**(I and J)** RACK1 MARylation-mediated G3BP1 localization to stress granules is dependent on the levels of stalled polysomes. Immunofluorescent staining assays of OVCAR3 cells with Dox-induced knockdown of endogenous RACK1 and re-expression of exogenous RACK1 subjected to 15 minutes of treatment with 250 μM sodium arsenite ($NaAsO_2$) (*left*, "Untreated"). The cells were also treated with 10 μg/mL puromycin for (I) 15 minutes or (J) 30 minutes prior to 15 minutes of treatment with 250 μM sodium arsenite ($NaAsO_2$) (*right*, "Puromycin"). Staining for HA (HA-RACK1) and G3BP1. DNA was stained with DAPI. Scale bar is 15 μm.
**(K and L)** Quantification of multiple experiments like those shown in (K) panel I above and (L) panel J above. Each bar represents the mean + SEM of distinct G3BP1 foci (n = 3, One-way ANOVA. * $p < 0.05$ and ns not significant).
**(M and N)** RACK1-Mut expression does not alter global protein synthesis under stress in OVCAR3 cells. (M) Immunoblot analysis of puromycin incorporation assays from OVCAR3 cells subjected to Dox-induced knockdown of endogenous and re-expression of RACK1 followed by 15 minutes of treatment with 250 μM sodium arsenite. β-tubulin serves as a loading control. (N) Quantification of immunoblot experiments like those shown in (M). Each bar in the graph represents the mean + SEM of the relative levels of puromycin incorporation (n = 3, One-way ANOVA, ns not significant).

### Supplemental Figure S1.

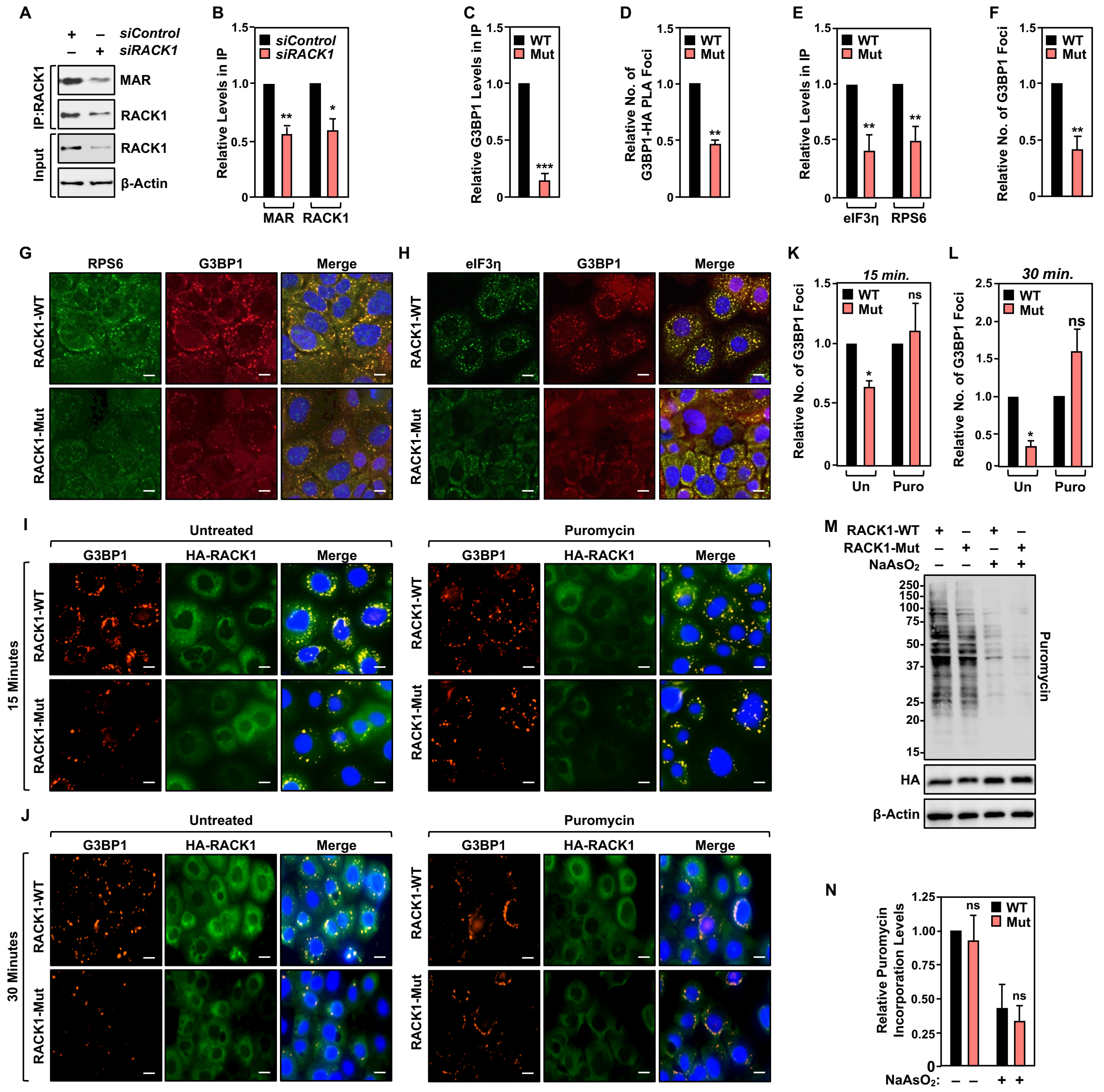

*<Fig. S2 is on the next page>*

**Supplemental Figure S2. PARP14 mediates RACK1 MARylation.**

**(A)** OVCAR3 cells were subjected to knockdown with two different siRNAs targeting each of the expressed cytosolic MARTs. Representative images from PLAs using MAR and RACK1 antibodies. DNA was stained with DAPI. Scale bar is 15 µm.

**(B)** Quantification of multiple experiments like the one shown in Figure 3B. Each bar represents the mean + SEM of MAR-RACK1 PLA foci ($n = 3$, Student's t-test, *** $p < 0.001$).

**(C and D)** PARP14 inhibition reduces G3BP1 interaction with translation factors that are key components of stress granules. Immunofluorescent staining assays of OVCAR3 cells treated with 10 µM PARP14 inhibitor (PARP14i) for 24 hours and subjected to 15 minutes of treatment with 250 µM sodium arsenite ($NaAsO_2$). Staining for (B) RPS6 and G3BP1, (C) eIF3η and G3BP1. DNA was stained with DAPI. Scale bar is 15 µm.

**(E and F)** PARP14 inhibition blocks RACK1 MARylation in ovarian cancer cells. PLA using MAR and RACK1 antibodies in (E) SKOV3 cells and (F) HCC5044 cells treated with 10 µM PARP14 inhibitor (PARP14i) for 24 hours and subjected to 15 minutes of treatment with 250 µM sodium arsenite ($NaAsO_2$). DNA was stained with DAPI. Scale bar is 15 µm.

**(G and H)** PARP14 inhibition blocks the assembly of G3BP1-containing stress granules in ovarian cancer cells. Immunofluorescent staining assays in (G) SKOV3 and (H) HCC5044 cells treated with 10 µM PARP14 inhibitor (PARP14i) for 24 hours and subjected to 15 minutes of treatment with 250 µM sodium arsenite ($NaAsO_2$). DNA was stained with DAPI. Scale bar is 15 µm.

#### Supplemental Figure S2.

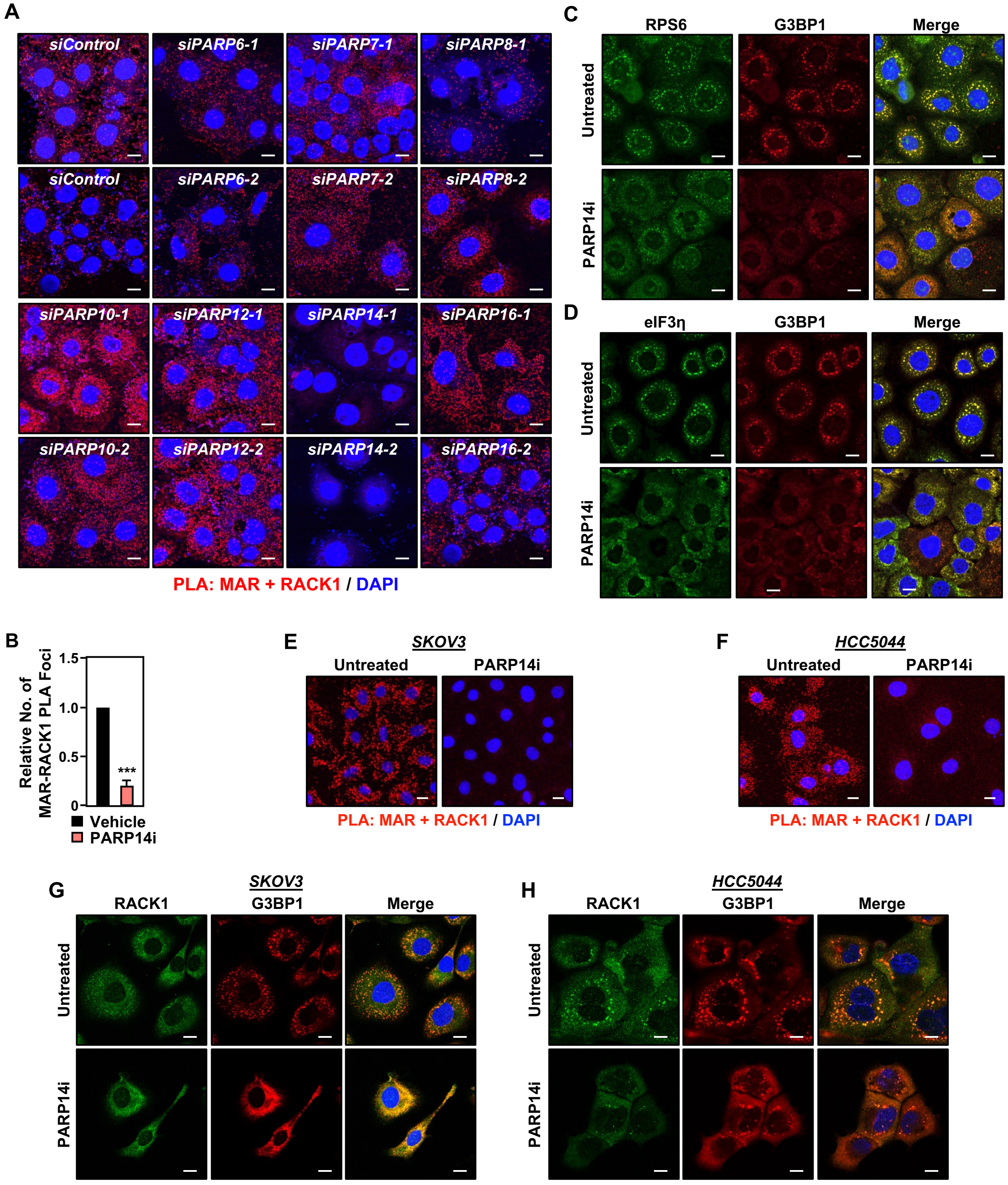

*<Fig. S3 is on the next page>*

**Supplemental Figure S3. PARP14 inhibition sensitizes ovarian cancer cells to stress and inhibits their growth.**
**(A and B)** Growth curves in the presence or absence of 10 μM PARP14 inhibitor (PARP14i) and 3 nM thapsigargin (Thps) for the indicated times. (A) SKOV3 cells and (B) HCC5044 cells. Each point represents the mean ± SEM of the growth of the cells relative to Day 0 of treatments (n = 3, two-way ANOVA, ** $p < 0.001$).
**(C)** RACK1-Mut expressing cells are sensitive to ER stress, which inhibits their growth. Growth curves of OVCAR3 cells with Dox-induced knockdown of endogenous RACK1 and re-expression of exogenous RACK1 (WT or Mut) in the presence or absence of 5 μM carboplatin for the indicated times. Each point represents the mean ± SEM of the growth of the cells relative to Day 0 of treatment (n = 3, two-way ANOVA, * $p < 0.05$).
**(D)** PARP14 inhibition sensitizes ovarian cancer cells to ER stress and inhibits their growth. Growth curves of OVCAR3 cells in the presence or absence of 10 μM PARP14 inhibitor (PARP14i) and 5 μM carboplatin for the indicated times. Each point represents the mean ± SEM of the growth of the cells relative to Day 0 of treatment (n = 3, two-way ANOVA, * $p < 0.05$).
**(E through G)** Growth curves of OVCAR3 xenograft tumors in immunocompromised NSG mice. The xenograft tumors were established in immunocompromised NSG mice subjected to the experimental conditions and treatments indicated and grown until the mice reached the end-point for euthanasia as required by IACUC. (E) Growth of OVCAR3 xenograft tumors with Dox-induced knockdown of endogenous RACK1 and re-expression of exogenous RACK1 (WT or Mut) for the indicated times. n = 10 or 8 mice (WT or Mut, respectively), Student's t-test, * $p < 0.05$; ** $p < 0.02$. (F) Immunoblot analysis of HA-RACK (WT or Mut) expression in the tumors at the end of the experiment. (G) Growth of OVCAR3 xenograft tumors with or without PARP14 inhibitor (PARP14i) treatment for the indicated times. n = 5 or 6 mice (vehicle or PARP14i, respectively), Student's t-test, * $p < 0.05$.
**(H)** RACK1-Mut expressing OVCAR3 cells exhibit greater ER stress. Immunoblot analysis of lysates from OVCAR3 cells with Dox-induced knockdown of endogenous RACK1 and re-expression of exogenous RACK1 (WT or Mut) the presence or absence of 3 nM thapsigargin (Thps) for 24 hours as indicated. Blotting for phospho- eIF2a (p-eIF2a), total eIF2a (eIF2a), cleaved caspase-3 (cl Caspase-3), HA, and β-tubulin (loading control) as indicated.
**(I through K)** PARP14 inhibitor (PARPi)-treated cells exhibit greater ER stress. Immunoblot analysis of lysates from (I) OVCAR3, (J) SKOV3, and (K) HCC5044 cells treated with 10 μM PARP14i and 3 nM thapsigargin for 24 hours. Blotting for phospho- eIF2a (p-eIF2a), total eIF2a (eIF2a), HA, and β-tubulin (loading control) as indicated.

**Supplemental Figure S3.**

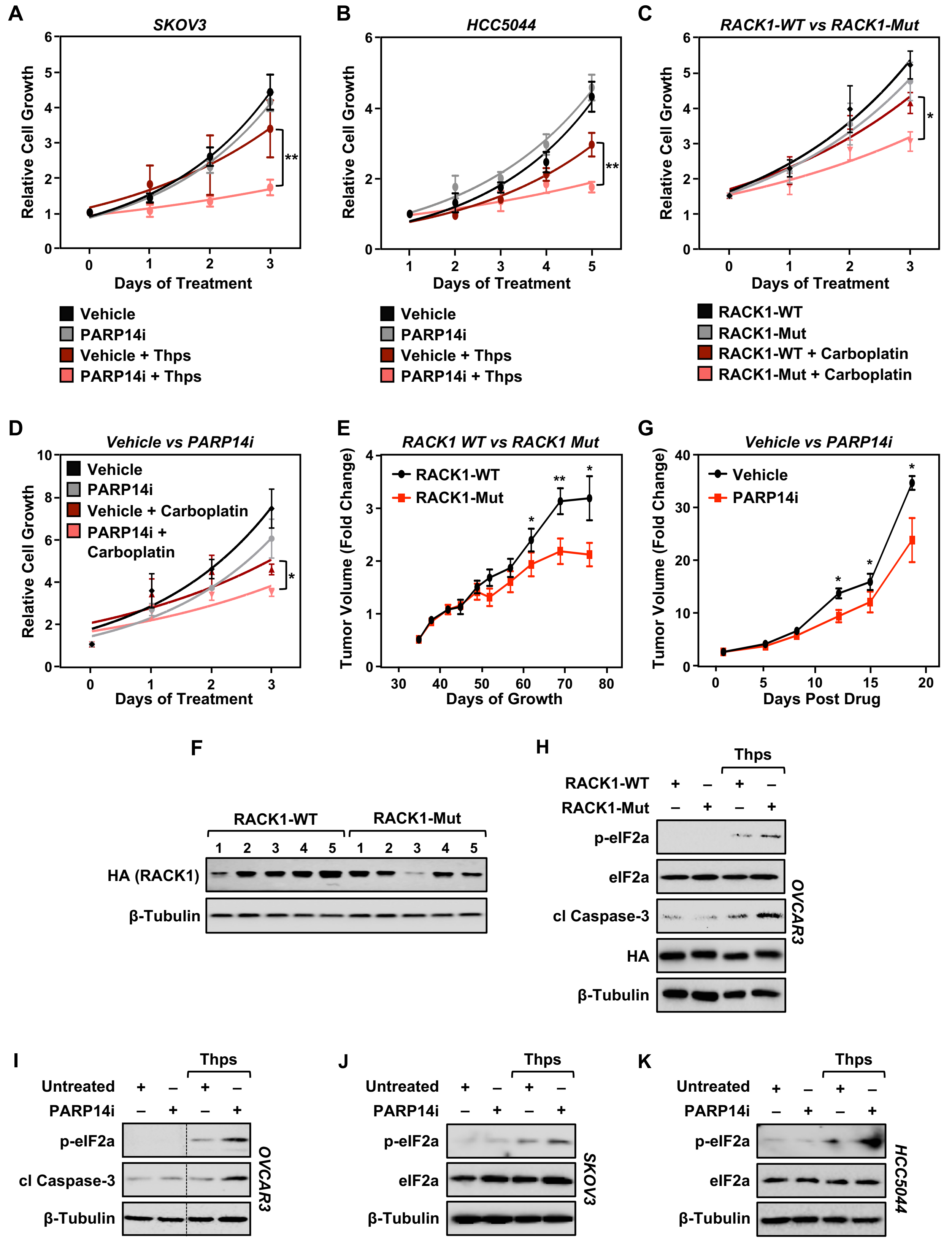

*<Fig. S4>*

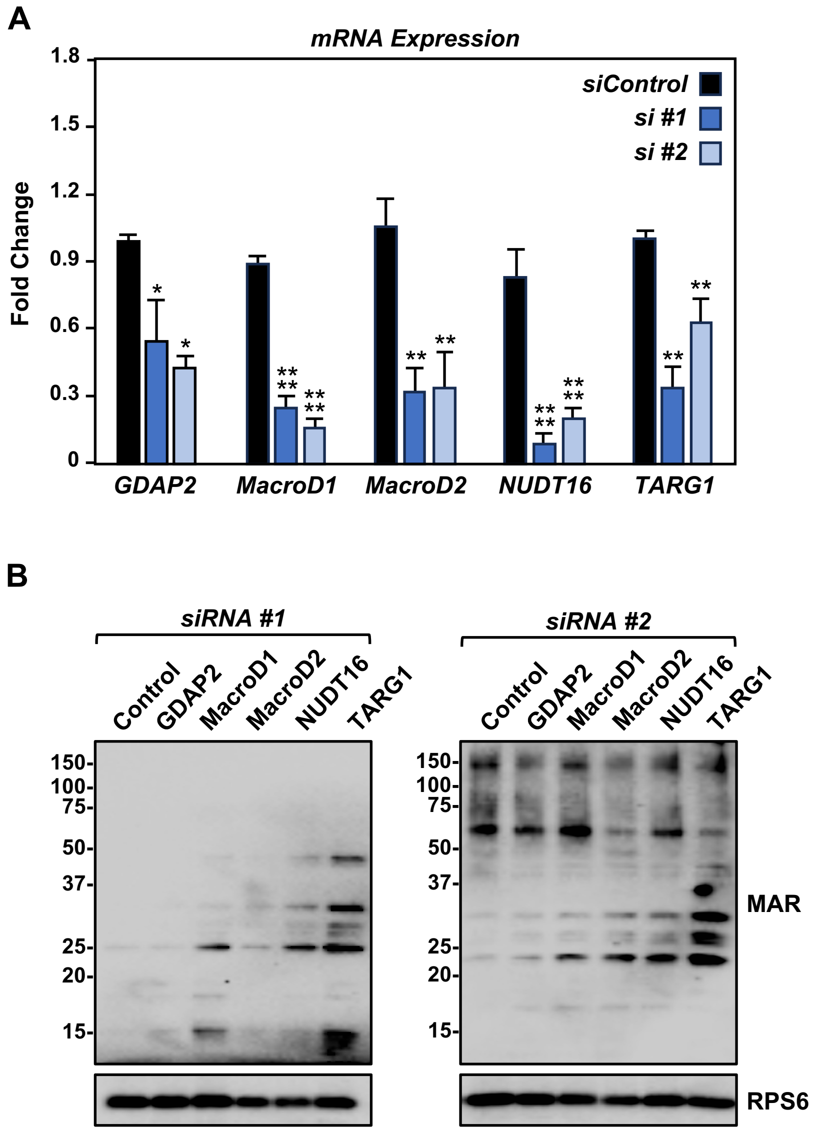


**Supplemental Figure S4. Small-scale siRNA screen to identify a ribosomal MAR hydrolase.** **(A)** HEK-293T cells were subjected to knockdown with two different siRNAs targeting each of the indicated ADPR hydrolases. RT-qPCR analysis of the mRNAs encoding the ADPR hydrolases. Each bar in the graph represents the mean + SEM of the mRNA levels of the indicated ADPR hydrolase, normalized to the levels of *GAPDH* mRNA (n = 3, pairwise t-tests with the Holm-Sidak correction, *p < 0.05, ** p < 0.001, **** p < 0.0001).
**(B)** Immunoblot analysis of ribosome MARylation from cells treated as described in (A). RPS6 is a loading control.

*<Fig. S5 is on the next page>*

**Supplemental Figure S5. Depletion of TARG1 enhances stress granule assembly by increasing RACK1 MARylation.**

**(A)** Quantification of multiple experiments like the one shown in Figure 5A. Each bar represents the mean + SEM of the level of G3BP1 in RACK1 immunoprecipitates (n = 4, One-way ANOVA, * $p < 0.05$).

**(B)** Quantification of multiple experiments like the one shown in Figure 5B. Each bar represents the mean + SEM of MAR-RACK1 PLA foci (n = 3, Student's t-test, *** $p < 0.001$).

**(C)** Quantification of multiple experiments like the one shown in Figure 5C. Each bar represents the mean + SEM of distinct G3BP1 foci (n = 3, Student's t-test, * $p < 0.05$, ** $p < 0.01$).

**(D and E)** siRNA-mediated *TARG1* depletion increases RACK1 MARylation in (D) SKOV3 cells and (E) HCC5044 cells subjected to 15 minutes of treatment with 250 μM sodium arsenite ($NaAsO_2$). PLA using MAR and RACK1 antibodies. DNA was stained with DAPI. Scale bar is 15 μm.

**(F and G)** siRNA-mediated *TARG1* depletion increases the assembly of G3BP1-containing stress granules. Immunofluorescent staining assays for RACK1 and G3BP1 in (F) SKOV3 cells and (G) HCC5044 cells with siRNA-mediated knockdown of *TARG1* subjected to 15 minutes of treatment with 250 μM sodium arsenite ($NaAsO_2$). DNA was stained with DAPI. Scale bar is 15 μm.

**(H)** Quantification of multiple experiments like the one shown in Figure 6B, top. Each bar represents the mean + SEM of TARG1-RACK1 PLA foci (n = 3, ANOVA, * $p < 0.05$, ** $p < 0.01$).

**(I)** Quantification of multiple experiments like the one shown in Figure 6B, bottom. Each bar represents the mean + SEM of MAR-RACK1 PLA foci (n = 3, ANOVA, * $p < 0.05$, ** $p < 0.01$).

### Supplemental Figure S5.

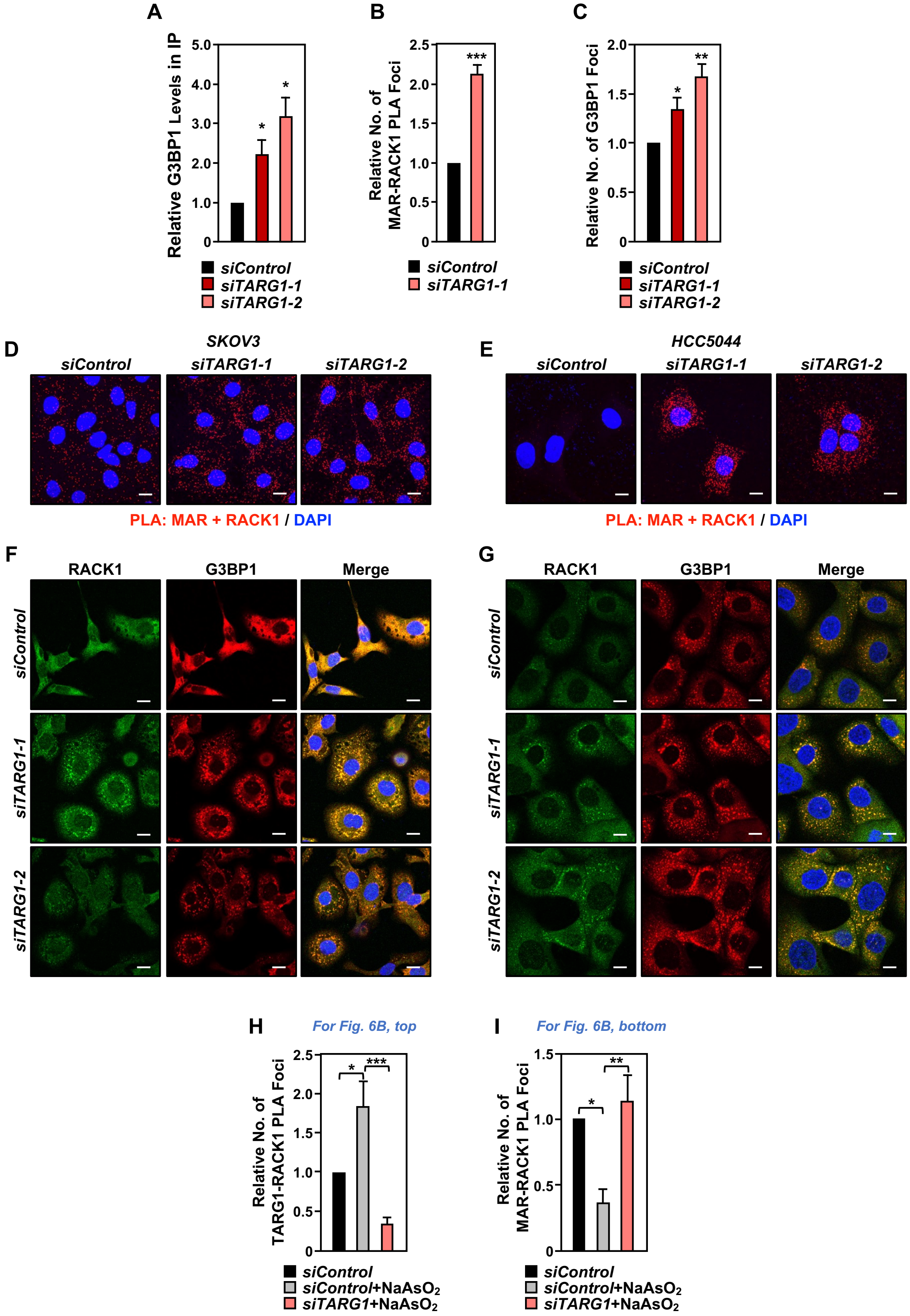
